# Structure of SWI/SNF chromatin remodeller RSC bound to a nucleosome

**DOI:** 10.1101/800508

**Authors:** Felix R. Wagner, Christian Dienemann, Haibo Wang, Alexandra Stützer, Dimitry Tegunov, Henning Urlaub, Patrick Cramer

## Abstract

Chromatin remodelling complexes of the SWI/SNF family function in the formation of nucleosome-depleted regions and transcriptionally active promoters in the eukaryote genome. The structure of the *Saccharomyces cerevisiae* SWI/SNF family member RSC in complex with a nucleosome substrate reveals five protein modules and suggests key features of the remodelling mechanism. A DNA-interacting module grasps extra-nucleosomal DNA and helps to recruit RSC to promoters. The ATPase and arm modules sandwich the nucleosome disc with their ‘SnAC’ and ‘finger’ elements, respectively. The translocase motor engages with the edge of the nucleosome at superhelical location +2 to pump DNA along the nucleosome, resulting in a sliding of the histone octamer along DNA. The results elucidate how nucleosome-depleted regions are formed and provide a basis for understanding human chromatin remodelling complexes of the SWI/SNF family and the consequences of cancer mutations that frequently occur in these complexes.

---

Nucleosomes that occupy gene promoters inhibit transcription initiation by RNA polymerase II and must be evicted or slid along DNA to establish nucleosome-depleted regions (NDRs) and to create active promoters^1^. Nucleosomes are evicted or slid by chromatin remodelling complexes that hydrolyse adenosine triphosphate (ATP)^2^. Remodelling complexes of the SWI/SNF family are of particular importance for NDR formation and transcription, and mutations in these complexes are linked to human cancers^3, 4^. The yeast *Saccharomyces cerevisiae* contains two complexes of this family, the SWI/SNF complex^5, 6^, and the essential and abundant 16-subunit complex RSC (‘Remodels the Structure of Chromatin’)^7^.

RSC contains the ATPase subunit Sth1 that functions as a DNA translocase^8–10^ and is required for normal transcription activity^11^. RSC can remove nucleosomes from promoters in reconstitution assays *in vitro*^12^. *In vivo,* RSC localizes to promoter regions^13^, and its loss leads to reoccupation of NDRs with nucleosomes^14^. RSC can bind and position the specialized +1 and −1 nucleosomes^15–17^ that flank NDRs on the downstream and upstream side, respectively^1, 4^. RSC can recognize poly(A) and GC-rich elements in promoter DNA^16, 18, 19^. The arrangement of these elements determines the strength and directionality of RSC action on promoter nucleosomes^20^.

Understanding how promoter nucleosomes are remodelled and how NDRs are established requires structural studies of RSC and its functional complexes. Electron microscopy (EM) studies of RSC showed a flexible structure with a central cavity that was suggested to bind a nucleosome^21–23^. However, these studies were limited to low resolution, which prevented molecular-mechanistic insights. Here we present the cryo-EM structure of RSC engaged with a nucleosome substrate. The results reveal the intricate subunit architecture of RSC, show how RSC engages with the nucleosome and adjacent DNA, and elucidate substrate recognition and remodelling mechanisms.

## Structure of RSC-nucleosome complex

Endogenous RSC was isolated from the yeast *Saccharomyces cerevisiae* via affinity purification of the tagged subunit Rsc2 (Extended Data Fig. 1a) (Methods). A RSC-nucleosome complex was assembled with DNA overhangs on each end of the nucleosome in the presence of the ATPase transition state analogue ADP-BeF_3_ (Extended Data Fig. 1b). Cryo-EM analysis resulted in a medium-resolution reconstruction that revealed the nucleosome, four turns of DNA exiting from one side of the nucleosome, and five RSC modules that we refer to as ATPase, ARP, body, arm, and DNA-interaction module (DIM) (Figure 1a; Extended Data Fig. 2). Focussed 3D classification enabled modelling of the nucleosome and associated ATPase with the use of a related structure^24^, and placement of an adapted ARP module structure^25^ (Extended Data Fig. 2). We also subjected the free RSC complex to cryo-EM analysis, and resolved the body and arm modules at resolutions of 3.6 Å and 3.8 Å, respectively (Extended Data Figs. 3, 4a-c). This led to a structural model of the RSC-nucleosome complex that only lacks the DIM module and agrees with lysine-lysine crosslinking information (Extended Data Fig. 1c, d).

**Figure 1.**
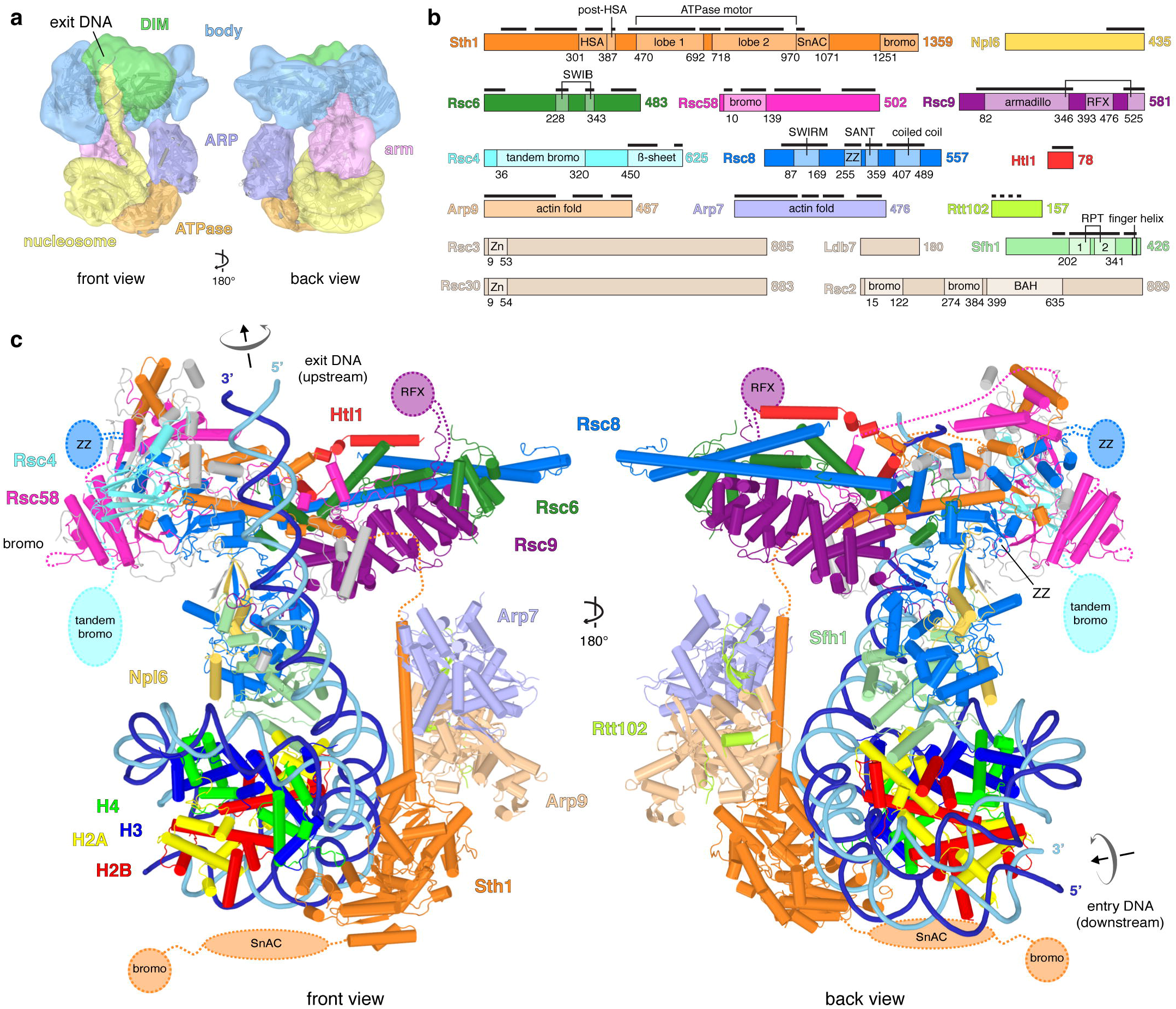
Structure of RSC-nucleosome complex. **a.** Overall architecture presented in two views of the low pass-filtered cryo-EM density. The five RSC modules are in different colours. Colour code for modules used throughout. The nucleosome substrate with exit DNA is in yellow. DIM, DNA-interaction module. **b.** Schematic of RSC subunit domain architecture. Colour code for subunits used throughout. Domain boundaries are marked with residue numbers and black bars indicate modelled regions. HSA, helicase-SANT-associated; SnAC, Snf2 ATP coupling; bromo, bromodomain; armadillo, armadillo repeat fold; RFX, DNA-binding RFX-type winged-helix; SWIRM, Swi3 Rsc8 Moira; ZZ, ZZ-type zinc finger; SANT, Swi3 Ada N-Cor TFIIIB; coiled coil, C-terminal helix forming coiled coil-like structure; Zn, Zn(2)-C6 fungal-type zinc finger; RPT, repeat; BAH, bromo-adjacent homology. **c.** Cartoon representation. Unassigned elements shown in grey. Mobile domains depicted schematically. Arrows indicate directionality of DNA translocation.

**Figure 2.**
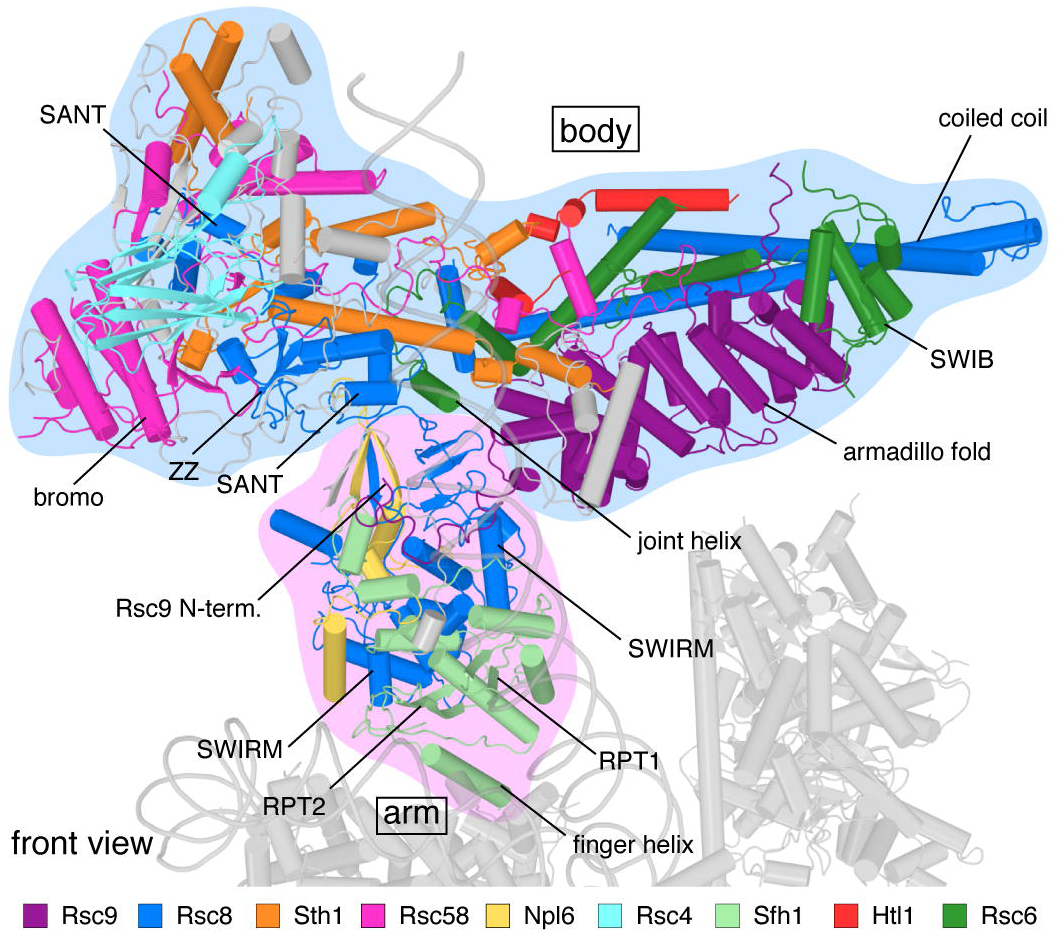
Structure of RSC body and arm modules. Cartoon representation viewed as in Figure 1. Important structural elements are labelled.

**Figure 3.**
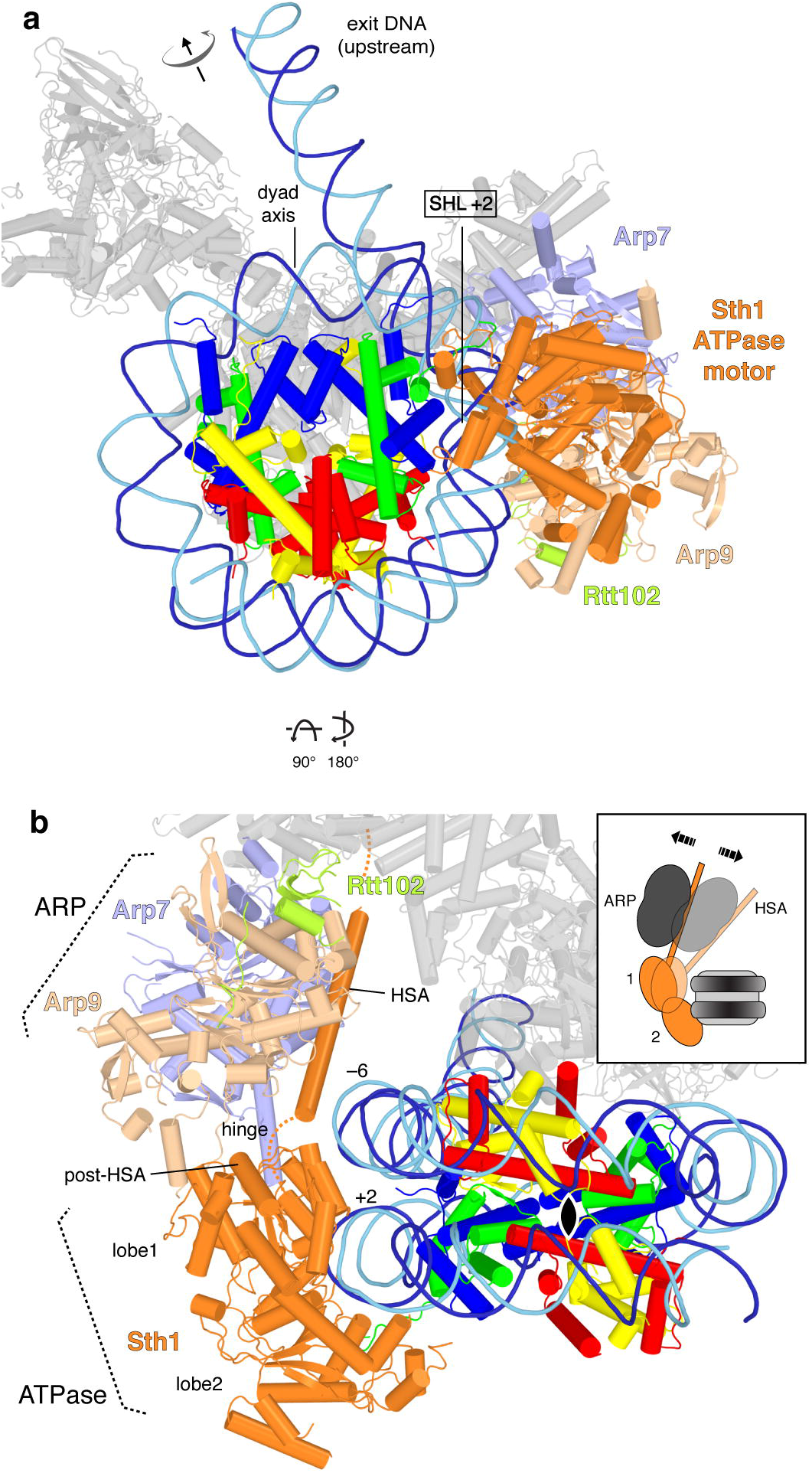
ATPase-nucleosome interactions. **a.** Contacts of Sth1 ATPase motor (orange) with the nucleosome. View as in Fig. 1c, left, but rotated by 45° around a horizontal axis. Arrows indicate directionality of DNA translocation. **b.** View along the nucleosome dyad (black oval). View is as in Fig. 1c, right, but rotated by 45° around a horizontal axis.

**Figure 4.**
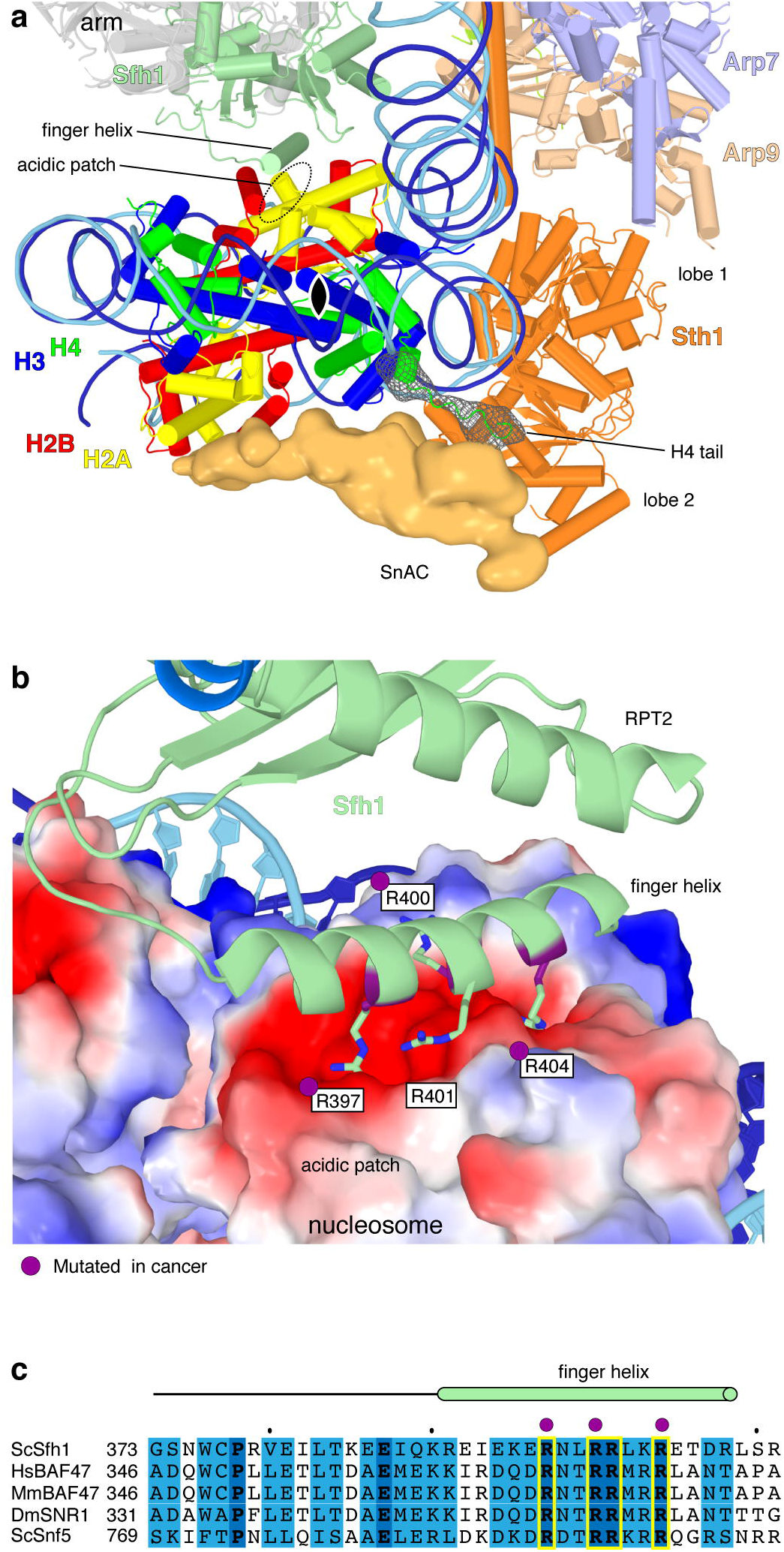
RSC sandwiches the nucleosome. **a.** RSC-nucleosome interactions viewed along the nucleosome dyad (black oval). On the outer face of the histone octamer, densities for the Sth1 SnAC domain and the histone H4 tail are shown as an orange surface and a green mesh, respectively. On the inner face, the arm module and Sfh1 finger helix are depicted. **b.** Interaction of the Sfh1 finger helix with the acidic patch of the inner face of the histone octamer (surface representation coloured by electrostatic charge; red, negative; blue, positive). Conserved arginine residues are shown with side chains. Residues mutated in human cancer (Methods) are highlighted in purple. **c.** Sequence alignment of the finger helix region (red cylinder) in *S. cerevisiae* (Sc) Sfh1 with its homologs *H. sapiens* (Hs) BAF47, *M. musculus* (Mm) BAF47, *D. melanogaster* (Dm) SNR1 and ScSnf5. Invariant and conserved residues in dark and light blue, respectively. Arginine residues shown in (b) highlighted in yellow. A purple dot marks residues mutated in cancer (Methods).

The structure reveals the intricate architecture of RSC (Figure 1, **Supplementary Video 1**). The body module contains subunits Rsc4, Rsc6, Rsc8, Rsc9, Rsc58, Htl1, and the N-terminal region of Sth1 (Figure 2, Extended Data Table 1). The ARP module is flexibly tethered to the body and comprises the helicase-SANT associated (HSA) region of Sth1, the actin-related proteins Arp7 and Arp9, and subunit Rtt102. The C-terminal region of Sth1 extends from the HSA region and forms the ATPase module (Extended Data Fig. 5a). The arm module protrudes from the body and contains subunit Sfh1 and parts of Rsc8, Npl6, and Rsc9 (Figure 2). The arm and body modules are tightly connected by two copies of Rsc8 that adopt different structures (Extended Data Fig. 5b). The N-terminal SWIRM domains of Rsc8 reside in the arm, whereas the SANT domains and one of the ZZ zinc finger domains reside in the body, as do the long C-terminal helices. The RSC structure and observed subunit interactions explain the requirement of the Rsc4 C-terminal region for cell growth^26^, the known interaction between Rsc6 and Rsc8^27^, and lethal effects of Rsc58 truncation^28^.

**Figure 5.**
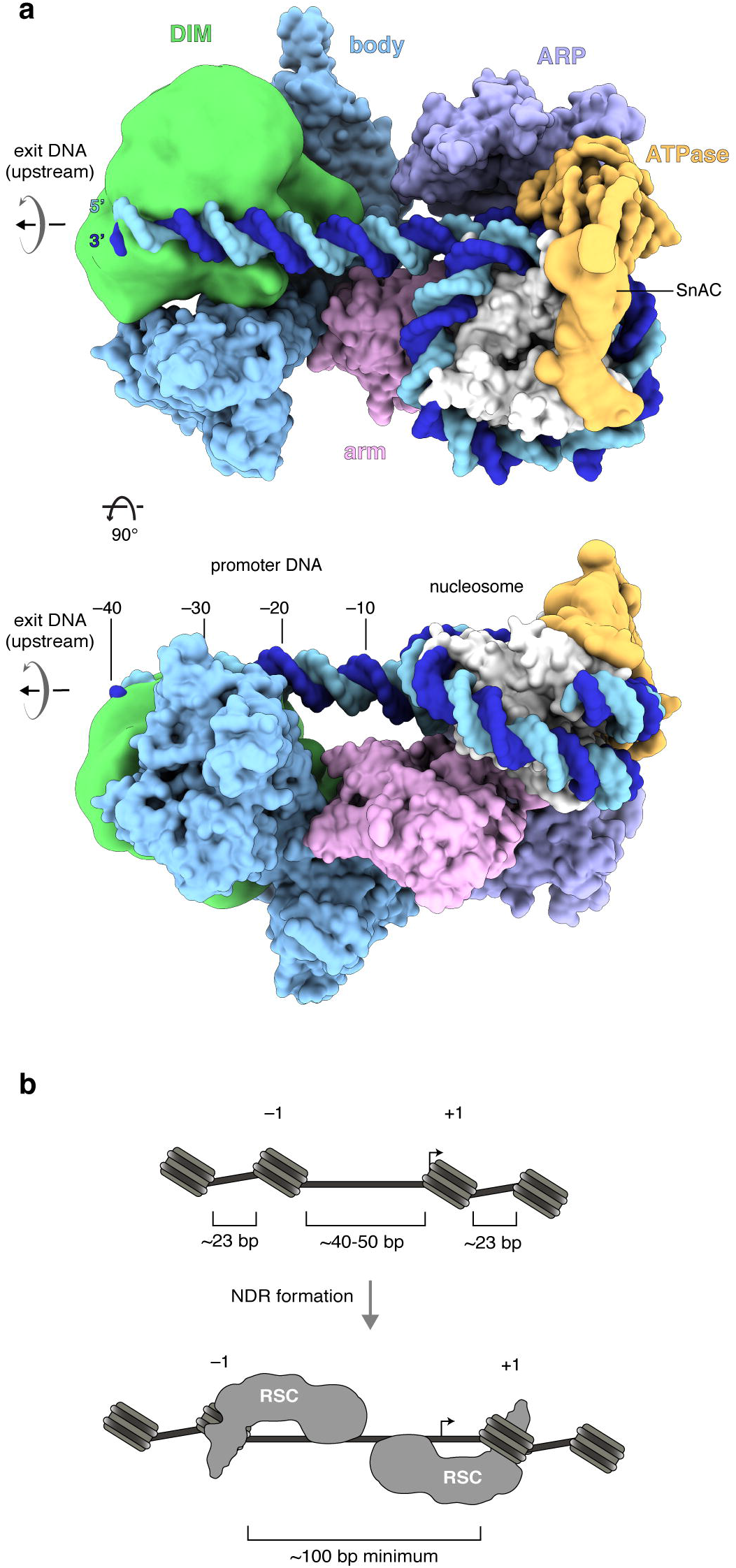
DNA recognition and NDR formation. **a.** Space-filling RSC-nucleosome structure with DIM (green) and SnAC (orange) densities. View on the top as in Fig. 1c, left, but rotated by 90°. Arrows indicate directionality of DNA translocation. Number of upstream DNA base pairs relative to SHL –7 is provided. **b.** Schematic of a promoter before (top) and after (bottom) RSC remodelling shows NDR formation by sliding the flanking –1 and +1 nucleosomes away from the NDR center. Arrows indicate the transcription start site.

RSC also contains six domains that are implicated in interactions with histone tails. The N-terminal bromodomain in Rsc58 locates to the surface of the body (Extended Data Fig. 5c). The five other domains are mobile, and include a bromodomain in Sth1, two bromodomains in Rsc2, a BAH domain in Rsc2 that binds histone H3^29^, and a tandem bromodomain in Rsc4 that interacts with acetylated H3 tails^26^, in particular acetylated lysine K14^26, 30^. RSC also contains five putative DNA-binding domains, of which four are mobile. These include the zinc finger domains in subunits Rsc3 and Rsc30, an RFX domain in subunit Rsc9, and a ZZ finger domain in one of the two Rsc8 subunits. In summary, RSC consists of five modules and nine flexibly connected domains, of which some are suggested to be involved in substrate selection via the recognition of histone modifications and DNA sequence features.

## ATPase binding and translocation

RSC engages in multivalent interactions with its substrate, contacting both DNA and histones (Figure 1). The ATPase and arm modules interact with the nucleosome, whereas the DIM module engages with DNA exiting from the nucleosome. The ATPase module binds the edge of the nucleosome, contacting both DNA gyres in a conformation poised for translocation activity (Figure 3a). The two lobes of the ATPase motor domain contact one gyre at superhelical location (SHL) +2 and adopt the same relative orientation as in the structure of the related SWI/SNF ATPase Snf2 bound to a nucleosome^24^. The N-terminal ATPase lobe 1 also binds the second DNA gyre around SHL –6 (Figure 3b), a location where the N-terminal tail of histone H3 is expected to protrude (Extended Data Figure 4d). Considering the known directionality of the translocase^31^, we arrive at the model that the RSC ATPase motor pumps DNA towards the nucleosome dyad and along the octamer surface in the exit direction, which corresponds to the upstream direction of transcription, thus liberating more promoter DNA.

The ARP module couples RSC ATPase activity to DNA translocation and regulates the remodelling activity^9, 25, 32^. Our results suggest that this regulation involves changes in the position of the mobile ARP module that influence the conformation and mobility of the ATPase lobe 1 and its interactions with both DNA gyres (Figure 3b). These changes are likely transmitted through the hinge region between the HSA region and lobe 1 that includes the ‘post-HSA’ region of Sth1. Mutations of the post-HSA region increase ATPase activity and DNA translocation, suggesting that the hinge acts as a throttle for the ATPase^8–10^. The ARP module adopts a defined position in the RSC-nucleosome complex, but it is mobile in the free RSC structure. Based on these results, we propose that the position of the mobile ARP module can influence the conformation and motility of the bilobal ATPase motor and thereby control the translocation activity of RSC (Figure 3b).

## Nucleosome sandwiching and sliding

The structure also suggests a model for how RSC can slide nucleosomes along DNA. RSC contacts the nucleosome disc not only at the edge, but also binds both of its faces. The SnAC domain in subunit Sth1 binds the outer face of the histone octamer, whereas the arm module binds the inner face (Figure 4a). Sandwiching interactions would retain the histone octamer and enable the ATPase motor to pump DNA around it, effectively sliding the octamer downstream on DNA. Consistent with this model, the SnAC domain in the SWI/SNF homologue Snf2 is important for remodelling *in vivo* and biochemical data suggested that it acts as a histone anchor that is required for nucleosome sliding^33^. The strength of the SnAC-histone octamer contact may be influenced by the N-terminal tail of histone H4, which binds at the interface of the SnAC and ATPase motor of Sth1 (Figure 4a). Since histone acetylation can impair octamer transfer by RSC to the histone chaperone Nap1^34^, this leads to the intriguing model that histone acetylation may strengthen the sandwiching contacts, thereby impairing octamer eviction and favouring nucleosome sliding.

Binding of the arm module to the inner face of the histone octamer is mediated by an exposed ‘finger’ helix, which resides in the C-terminal region of subunit Sfh1 that is required for normal cell growth^35^ (Figure 4a, Extended Data Figs. 1d, 4e). The finger helix contains four arginine residues (R397, R400, R401 and R404) that contact the acidic patch of the octamer. Three of these arginines are known to be mutated in human cancers (Figure 4b), pointing to the functional significance of the finger helix-acidic patch interaction. The finger helix and its arginine residues are highly conserved in Sfh1 homologs throughout eukaryotes (Figure 4c). The SnAC domain is also conserved over species and between SWI/SNF complexes^36^, suggesting that the sandwiching mechanism of nucleosome sliding is used by all SWI/SNF family complexes.

The arm module and its finger helix may also contribute to substrate selection. RSC preferentially recognizes nucleosomes that contain the histone variant H2A.Z^37^. Such nucleosomes show a more extended acidic patch^38^ and may have increased affinity for the basic RSC finger. The arm module may also contact the C-terminal tail of H2A.Z (Extended Data Fig. 4d) that differs in ten amino acid residues from the tail of H2A in yeast. The observed arm-octamer interaction also explains why ubiquitination of histone H2B counteracts RSC function^39^. The ubiquitin moiety attached to H2B residue K123 (human K120) is predicted to sterically interfere with the arm-octamer interaction (Extended Data Fig. 4f).

## DNA recognition and NDR formation

RSC not only binds the nucleosome, but also DNA that exits from it (Figure 5a). The DIM module contacts exiting DNA ∼20 – 40 bp upstream of SHL –7 of the nucleosome. This is in agreement with RSC protecting ∼50 bp of extra-nucleosomal DNA from nuclease digestion^15^. The DIM-DNA contact also explains how RSC recognizes specific DNA elements that are enriched in promoters^16, 18–20^. Consistent with crosslinking information, the RSC subunits Rsc2, Rsc3 and Rsc30 are located in the DIM (Extended Data Fig. 1d). Rsc3 and Rsc30 are known to interact^40^ and recognize a CGCG DNA element located upstream of the transcription start site^18^. They may bind DNA via their N-terminal zinc cluster domains^18, 40^. It remains to be seen to what extent promoter targeting by RSC depends on its binding to DNA sequence, histone modifications, and the presence of histone variant H2A.Z.

The results also elucidate the formation of NDRs. In *S. cerevisiae,* the DNA linker length between two nucleosomes is only ∼23 bp on average^41^. Steric considerations predict that RSC can enter chromatin only at sites where the length of the DNA linking two nucleosomes is at least 40 – 50 bp (Figure 5b). This can explain why RSC is targeted only to promoter regions, which are intrinsically nucleosome-depleted to some extent. NDR formation involves sliding of both flanking nucleosomes away from the NDR center^12^. Here we have interpreted the RSC-nucleosome structure to describe RSC action on the +1 nucleosome, but the structure can equally describe RSC action on the –1 nucleosome. In the latter case, DNA exits in downstream direction, rather than upstream, the ATPase engages with SHL –2, rather than SHL +2, and DNA translocation slides the nucleosome upstream, rather than downstream. Provided that RSC remains bound to both flanking nucleosomes after remodelling, a minimum NDR size of ∼100 bp would result (Figure 5b). However, larger NDRs can be formed when RSC evicts a nucleosome^16^.

## SWI/SNF family and cancer

The PBAF complex is the human counterpart of RSC and contains subunits homologous to Sth1, Rsc6, the Rsc8 dimer, Sfh1, Arp7 and Arp9 (Extended Data Table 1). In addition, the PBAF subunit BAF200^42^ contains an armadillo repeat fold^43^ and likely corresponds to Rsc9. Further, the BAF180 subunit comprises regions that resemble Rsc2 and Rsc4^44^. Only the small RSC subunits Rsc58, Rtt102 and Htl1 lack obvious counterparts. Therefore, the yeast RSC structure is a good model for human PBAF. Projection of the homologous regions onto the RSC structure reveals that the ATPase, ARP and arm modules are well conserved in PBAF, and that the body is at least partially conserved (Figure 6a). The DIM differs substantially in PBAF because human counterparts of subunits Rsc3 and Rsc30 are not known. However, PBAF subunits contain 12 putative DNA-binding domains that may mediate DNA recognition (Extended Data Table 1).

**Figure 6.**
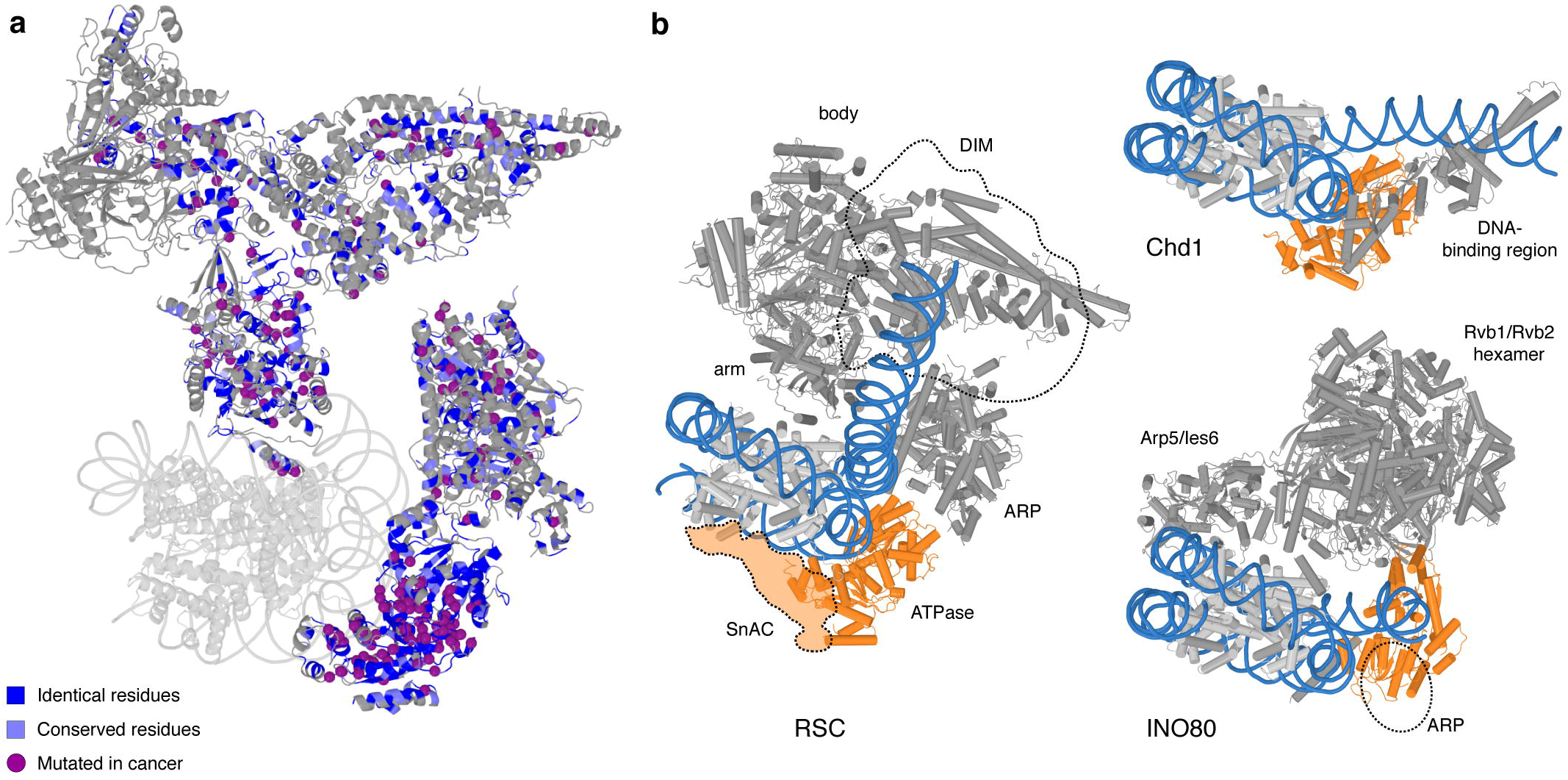
Remodeller families and cancer mutations. **a.** Conservation between SWI/SNF complexes RSC (yeast) and PBAF (human). Residues that are identical (blue) or conserved (light blue) in human PBAF highlighted on the RSC structure (grey). Purple spheres depict identical residues that are mutated in various cancers (Methods). **b.** Comparison of overall structure of RSC with complexes of CHD (yeast CHD1^47^) and INO80 (yeast INO80^51^) families. ATPase motor domains are shown in orange, DNA in blue.

The RSC structure also suggests the architecture of the related yeast SWI/SNF complex and its human counterpart BAF. Based on subunit composition and sequence homologies (Extended Data Figure 6), the yeast SWI/SNF complex contains RSC-related ATPase, ARP, and arm modules, whereas its body module is apparently smaller. The human BAF complex contains counterparts of RSC subunits Sth1, Sfh1, Arp7, Arp9, Rsc6, and Rsc8. The BAF subunit BAF250a is predicted to contain five armadillo repeats^45^, and is likely the counterpart of Rsc9. Thus, BAF also contains the ATPase, ARP and arm modules, and a body module that is at least partially conserved.

Due to these homologies, the RSC structure can be used to locate protein sites in PBAF that are known to be mutated in human cancers. This analysis shows that cancer-associated mutations are scattered throughout the remodelling complex (Figure 6a). Most mapped mutations are located inside the structured modules and are predicted to destabilize protein folds. However, mutations are particularly enriched within the ATPase, ARP and arm modules that surround and contact the nucleosome, suggesting that they cause functional defects.

## Diversity in chromatin remodellers

Finally, we compared the RSC-nucleosome structure with nucleosome complex structures of the three other families of chromatin remodelling factors (Figure 6b). Whereas RSC is a good model for the SWI/SNF family, factors of the ISWI, CHD and INO80 families are clearly distinct. With respect to the ISWI family, the ATPase motor binds SHL +2^46^, but other interactions have not been structurally resolved. With respect to the CHD family, the ATPase motor of yeast Chd1 also binds SHL +2, but its DNA-binding region engages with exiting DNA in close proximity to the nucleosome, leading to a different trajectory of exit DNA^47, 48^.

With regard to the INO80 family, the ATPase motor of the SWR1 complex also binds SHL +2^49^, whereas the ATPase of the INO80 complex binds SHL –6^50, 51^. INO80 also contains an ARP module^52^, which however contacts exit DNA in a manner that is distinct from the DIM-DNA contacts observed for RSC. The INO80 complex also contains flexible protein extensions, called the ‘Arp5 grappler’ and the ‘Ies RAR’, which can contact both faces of the histone octamer^51^, but these contacts differ substantially from the sandwiching interactions formed by RSC. In conclusion, the RSC-nucleosome structure provides mechanistic insights into the fourth family of chromatin remodelling complexes, and provides a basis for investigating the targeting, regulation, and cellular roles of SWI/SNF family complexes.

## Acknowledgements

We thank current and former members of the Cramer Laboratory, including S. Osman, G. Kokic, P. Seweryn, S, Schilbach, S. Neyer, and H. Hillen. F.R.W. was supported by a Boehringer Ingelheim Fonds PhD fellowship. H.U. was supported by the Deutsche Forschungsgemeinschaft (SFB860). P.C. was supported by the Deutsche Forschungsgemeinschaft (SFB860, SPP1935, EXC 2067/1-390729940), the European Research Council Advanced Investigator Grant TRANSREGULON (grant agreement No 693023), and the Volkswagen Foundation.

## Author contributions

F.R.W. carried out all experiments and data analysis unless stated otherwise. C.D. assisted with data collection and model building. A.S. and H.U. carried out crosslinking and mass spectrometry analysis. H.W. helped with nucleosome biochemistry. D.T. helped with cryo-EM data processing. P.C. designed and supervised the project. F.W. and P.C. wrote the manuscript, with input from all authors.

## Author information

The author declare that they have no competing financial interest. Correspondence and request of materials should be addressed to P.C.

## METHODS

### Preparation of RSC complex

The yeast *Saccharomyces cerevisiae* contains two isoforms of RSC that comprise either the subunit Rsc1 or its homologue Rsc2^53^ (Extended Data Table 1). We isolated the Rsc2-containing isoform. The *RSC2-TAP-HIS3* yeast strain (YSC1177-YLR357W) was purchased from the Dharmacon TAP-tagged open reading frame (ORF) library. A colony from a YP agar plate supplemented with 2% glucose (w/v) was used to prepare a 2 L pre-culture in YPD medium with 50 µg/mL ampicillin sodium salt and 12.5 µg/mL with OD_600_ of 1.6. Cells were fermented from OD_600_ ∼0.006 to OD_600_ ∼10 in 250 L of 3% YEP broth (w/v, Formedium) supplemented with 2% glucose, 50 g/L ampicillin sodium salt and 12.5 g/L tetracyclin-hydrochloride. The pellet was resuspended in cold 2x lysis buffer (100 mM HEPES pH 7.6, 20% glycerol (v/v), 1.4 M KAc, 2 mM MgCl_2_, 2 mM DTT and 3x protease inhibitor (100x: 0.028 mg/mL leupeptin, 0.137 mg/mL pepstatin A, 17 mg/mL PMSF, 33 mg/mL benzamidine)), frozen in liquid nitrogen to pea-sized granules and stored at –80 °C.

RSC was purified based on the TAP-tag purification strategy^8, 54, 55^, with several modifications. All purification procedures were performed at 4 °C unless stated otherwise. 600 g yeast granules were lysed by cryo-milling (Spex Freezer/Mill 6875D) and stored at –80 °C. Yeast powder was thawed at 30 °C, diluted with 100 mL 1x lysis buffer and cleared by centrifugation (25,200 xg). The supernatant was incubated for 6 h with 10 mL IgG Sepharose 6 Fast Flow resin (GE Healthcare) pre-equilibrated in lysis buffer. The resin was recovered by centrifugation (3,200 xg) and washed with 100 mL elution buffer A (50 mM K-HEPES, pH 7.6, 150 mM KAc, 10% glycerol (v/v), 3 mM CaCl_2_, 1 mM imidazole, 1 mM DTT, 0.5x protease inhibitor). IgG resin was resuspended in 10 mL elution buffer A, mixed with 2 mL calmodulin resin (Agilent Technologies) pre-equilibrated in elution buffer A and supplied with catalytic amounts of TEV protease. The resin was washed with 100 mL elution buffer A without protease inhibitors, and protein was eluted with 50 mL elution buffer B (3 mM EGTA instead of 3 mM CaCl_2_). Elution was applied to a HiTrap Q 1 mL HP column (GE Healthcare) pre-equilibrated with Q-150 buffer (50 mM HEPES pH 7.6, 150 mM KAc, 10% glycerol, 1 mM DTT) and washed with 10 CV Q-150 buffer. Protein was eluted with a linear gradient from 0 – 100 % buffer Q-1500 (1.5 M KAc instead of 150 mM KAc) over 50 CV. RSC-containing fractions were concentrated, dialysed overnight to 50 mM HEPES pH 7.6, 150 mM KAc, 10% glycerol, 5 mM MgCl_2_,1 mM DTT, and immediately used for cryo-EM sample preparation. Typical yields were 0.2 – 0.3 mg from 300 g yeast pellet.

### Preparation of nucleosome substrates

*Xenopus leavis* histones were expressed and purified as described^56, 57^. Briefly, histones were purified as inclusion bodies using a Dounce tissue grinder (Sigma-Aldrich). Histones were aliquoted, flash-frozen, lyophilised, and stored at –80 °C. For octamer preparation, lyophilised histones were resuspended in unfolding buffer (20 mM HEPES pH 7.5, 7 M guanidinium hydrochloride, 10 mM DTT) to a concentration of 3 mg/mL. Histones H2A, H2B, H3 and H4 were combined at a molar ratio of 1.2:1.2:1:1 and dialysed against two times 2 L of refolding buffer (10 mM HEPES pH 7.5, 2 M NaCl, 1 mM EDTA, 2.5 mM DTT) for a total of 12 h at 4 °C. The sample was concentrated and applied to a Superdex 200 Increase 10/300 size exclusion column pre-equilibrated with refolding buffer. Peak fractions were pooled and frozen in liquid nitrogen at a concentration of 1.34 mg/mL.

DNA fragments for nucleosome reconstruction were prepared by PCR as described^58^. gBlock DNA (IDT) containing the 145-bp Widom 601 sequence^59^ with a 55 bp extension at the 5’-end and a 37 bp extension at the 3’-end was used as a template together with two primers (forward: TCATTACCCAGCCCGCCTAG, reverse: CCTACGGACCGGATATCTTCCCTG). Reactions were pooled (42 mL) and DNA products recovered by phenol-chloroform-extraction. DNA was resuspended in MilliQ water and applied to a Superose 6 Increase 10/300 size exclusion chromatography column pre-equilibrated in gel filtration buffer (20 mM HEPES pH 7.5, 200 mM NaCl, 1 mM EDTA). Peak fractions were pooled, concentrated ten times, and stored at –20 °C.

Nucleosome reconstitution was performed as described^57^, with minor modifications. DNA and histone octamer were mixed at a 1:1.2 molar ratio in reconstitution buffer (20 mM HEPES pH 7.5, 1 mM EDTA, 2 mM DTT) containing 2 M NaCl and incubated for 30 min on ice. Sample was transferred to a Slide-A-Lyzer 3.5K MWCO MINI device and gradient-dialysed from 500 ml high salt reconstitution buffer against 2 L of low salt reconstitution buffer (20 mM NaCl) for 22 h. After a heat shift for 30 min at 50 °C, the sample was recovered and immediately used for complex formation.

### RSC-nucleosome complex formation

Newly prepared RSC complex was mixed with ADP-BeF_3_ at a final concentration of 1 mM and incubated on ice for 30 min. A 1.6-fold molar excess of the nucleosome substrate was added, the mixture incubated for 15 min at 30 °C and transferred back on ice. RSC-nucleosome complex was cross-linked using the GraFix method^60^. The sample was applied to a gradient generated from a 10% sucrose light solution (10% sucrose (w/v), 50 mM HEPES pH 7.6, 150 mM KAc, 5% glycerol (v/v), 2 mM MgCl_2_, 1 mM DTT, 0.5 mM ADP-BeF_3_) and a 25% sucrose heavy solution (25% sucrose (w/v) instead of 10%) containing 0.2% glutaraldehyde crosslinker with a BioComp Gradient Master 108 (BioComp Instruments). Centrifugation was carried out for 16 h at 32,000 rpm in a SW 60 Ti swinging-bucket rotor (Beckmann) at 4 °C. 200 µL fractions were collected and quenched with aspartate (pH 7.5) at a final concentration of 50 mM. Fractions containing RSC-nucleosome complex were dialysed for 8 h at 4 °C to 20 mM HEPES pH 7.6, 150 mM KAc, 1% glycerol (v/v), 3 mM MgCl_2_, 1 mM DTT and applied to cryo-EM grids.

### Cryo-EM analysis of RSC-nucleosome complex

RSC-nucleosome complex was absorbed to a thin carbon film before plunge freezing as described^61^, with minor modifications. A small, thin (∼3.1 nm) carbon film was floated from the mica sheet onto a 50 µL drop of sample and incubated for 2 – 3 min. The carbon film was recovered with copper R2/1 or R3.5/1 grids (Quantifoil) and vitrified by plunge-freezing in liquid ethane using a Vitrobot Mark IV (FEI) operated at 4 °C and 100% humidity.

Electron micrographs were acquired on an FEI Titan Krios G2 transmission election microscope operated at 300 keV in EFTEM mode, equipped with a Quantum LS 967 energy filter (Gatan), zero loss mode, 30 eV slit width, and a K2 Summit direct electron detector (Gatan) in counting mode. Automated data acquisition was done using the FEI EPU software package at a nominal magnification of 130,000x, resulting in a calibrated pixel size of 1.05 Å/px. Micrographs for the two datasets were collected at a dose rate of 4.78 e^−^/Å^2^/s over 10 s resulting in a total dose of 47.8 e^−^/Å^2^, and at a dose rate of 5.67 e^−^/Å^2^/s over 8 s resulting in a total dose of 45.4 e^−^/Å^2^, respectively. Both datasets were dose fractionated over 40 frames.

Dose weighting, CTF estimation and motion correction were carried out during data collection using Warp^62^. Automated particle picking by Warp resulted in 112,657 particles from the first dataset (4404 micrographs) and 1,119,875 particles from the second dataset (19,415 micrographs). Particle coordinates were exported, combined, extracted and processed using RELION 3.0^63^. Removal of bad particles through global 3D classifications with a negative stain reconstruction of the RSC complex as reference resulted in high-quality particles that could be refined to an overall map of the RSC remodeller together with the nucleosome (map 1) at a resolution of ∼15 Å. Further processing of the particles revealed great flexibility and dynamics which could not be resolved by focused 3D classifications and refinements.

The particles corresponding to the RSC-nucleosome map were reextracted centred on the nucleosome with a box mainly including the nucleosome and the ATPase module. Global 3D classification resulted in a good class that revealed the Sth1 subunit bound to the nucleosome. Focused 3D refinement excluding the Sth1 density provided a nucleosome map (map 2) at a resolution of 3.6 Å (gold-standard Fourier shell correlation 0.143 criterion) and a B-factor of –155 Å^2^. Improvement of the Sth1 density turned out to be very difficult and showed its highly dynamic nature in this sample. A strategy of focused 3D classification without image alignment on the Sth1 part, followed by a global 3D refinement and additional focused 3D classification on the combined Sth1-nucleosome density led to the best results. A focused 3D classification and postprocessing with automatic B-factor determination in RELION resulted in an overall resolution of the Sth1-nuclesome map of 4.3 Å (FSC 0.143 criterion) and B-factor of –186 Å^2^ (map 3) (Extended Data Fig. 2). The nucleosome alone could be resolved to 3.6 Å (FSC 0.143 criterion) using a B-factor of –156 Å^2^ (map 2) (Extended Data Fig. 2). Final focused maps were combined using the Frankenmap tool distributed with Warp (map 7) (Extended Data Fig. 2). Masks encompassing the regions of interest were created with UCSF Chimera^64^ and RELION.

### Cryo-EM analysis of the free RSC complex

Freshly purified RSC complex was mixed with ADP-BeF_3_ to a final concentration of 1 mM and incubated for 15 min on ice. BS3 (bis(sulfosuccinimidyl(suberate))) cross-linker (Thermo Fischer Scientific) was added to a final concentration of 1 mM, incubated on ice for 30 min before quenching with Tris-HCl, pH 7.5, and ammonium bicarbonate at a final concentration of 100 mM and 20 mM, respectively. After size exclusion chromatography using a Sepharose 6 Increase 3.2/300 column (GE Healthcare) pre-equilibrated in gel filtration buffer (50 mM HEPES pH 7.6, 150 mM KAc, 4 mM MgCl_2_, 1 mM DTT), peak fractions were immediately applied to cryo-EM grids. 4 µL of sample were applied to glow-discharged (Pelco easiGlow) R2/2 gold grids (Quantifoil). Grids were blotted and vitrified as described above.

Cryo-EM data was collected as described above, with small modifications. The energy filter slit width was set to 20 eV. Micrographs for the two 0° tilt datasets were collected at a dose rate of 4.88 e^−^/Å/s for 8 s resulting in a total dose of 39 e^−^/Å^2^ and at a dose rate of 5.02 e^−^/Å^2^/s over 9 s resulting in a total dose of 45.2 e^−^/Å^2^, respectively, and fractionated over 40 frames. The third, 25° tilted dataset was acquired in 44 frames at a dose rate of 4.99 e^−^/Å^2^/s for 11 s resulting in a total dose of 54.9 e^−^/Å^2^.

Pre-processing and particle picking was carried out as described above and resulted in 205,990 particles from the first dataset (1787 micrographs), 170,028 particles from the second dataset (1216 micrographs) and 475,168 particles from the tilted dataset (3158 micrographs). Particles were processed with global 3D classifications using RELION-3^63^ and a negative stain reconstruction of the RSC complex as a first reference to obtain an improved initial reference. All 1,009,020 particles were newly extracted and bad particles were sorted out in multiple rounds of global 3D classifications in combination with global 3D refinements. The best resulting class was refined with a mask excluding the flexible DNA-interaction module (DIM). Particles corresponding to this reconstruction were subjected to CTF refinement and Bayesian polishing in RELION. Using focused 3D refinements, the maps for the arm module, and body2 and body1 submodules were further improved. Postprocessing with automatic B-factor determination in RELION resulted in overall resolutions of 3.8 Å, 3.6 Å and 3.6 Å, respectively, and B-factors of –136 Å^2^, –100 Å^2^ and –103 Å^2^, respectively (Extended Data Figure 3). Final focused maps were combined with Warp (Extended Data Figure 3).

### Structural modelling

The lower resolution cryo-EM map 1 of the RSC-nucleosome complex was used to align the individually generated cryo-EM maps 2 – 8. The combined cryo-EM map 7 was used for model building of the Sth1 subunit bound to the nucleosome. The final map was created with the local resolution tool from RELION and a B-factor of –150 Å^2^. The structure of the yeast Snf2 bound to the nucleosome in the ADP-BeF_3_ state (PDB code 5Z3U)^24^ was used as basis for modelling. Published data together with the close homology between Sth1 and Snf2 (Extended Data Fig. 6) suggest that Sth1 also binds at SHL +2. The remodeler and the nucleosome part were fitted separately. The *Xenopus laevis* histones and Widom 601 sequence of PDB 5Z3U were the same as used in our study. The nucleosome structure was rigid-body fitted into our cryo-EM map in UCSF Chimera^64^ and the entry side DNA and histone tails trimmed according to the density in COOT^65^. Due to lower resolution, amino acid side chains of residues 15 – 22 of H4 (chain B) were stubbed in COOT. The nucleosome structure was flexibly fitted using Namdinator^66^ and real space refined in PHENIX^67^ with secondary structure restraints (including base paring and base stacking restraints).

High conservation of amino acids between Sth1 and Snf2 (Extended Data Figure 6) allowed for generation of a Sth1 homology model with Rosetta^68, 69^. The homology model was trimmed according to the density in COOT, Brace-II helix was removed, and amino acid side chains were stubbed owing to the lower resolution of the map area before rigid-body docking using UCSF Chimera. Additional real space refinement with secondary structure restraints (including base paring and base stacking restraints) was performed in PHENIX. The overhanging exit side DNA was modelled by generating a bend B-DNA following the density in map 1 in 3D-DART^70^. The DNA duplex was connected to the nucleosomal Widom 601 DNA and geometry optimized with base pairing and base stacking restraints in PHENIX.

Map 1 allowed for the rigid-body docking of the crystal structure of the Arp module bound to the Snf2 HSA region (PDB code 4I6M)^25^ using UCSF Chimera. The amino acid residues of the Snf2 HSA helix were mutated to the ones from Sth1 according to sequence alignment (Extended Data Fig. 6) starting at the C-terminus and ignoring gaps. The model for the Sth1 HSA helix is thus an extrapolation based on the strong α-helical secondary structure prediction and the register might differ slightly^25^.

The combined cryo-EM map 8 and the focused refined maps 4 – 6 were also used for model building. SWISS-MODEL^43, 71^ was used to generate homology models for the Rsc58 N-terminal bromodomain (PDB code 3LJW)^72^, the Rsc6 SWIB domain (PDB code 1UHR), the Rsc8 SWIRM (PDB code 2FQ3)^73^, SANT (PDB code 2YUS) and ZZ zinc finger domains (PDB code 1TOT)^74^, the Rsc9 armadillo-like domain (PDB code 4V3Q)^75^ and the Sfh1 RPT1 and RPT2 domains (PDB code 6AX5). The homology models were rigid-body placed using UCSF Chimera^64^ and manually adjusted and re-build in COOT^65^.

The quality of the maps allowed for *de novo* building of the other model parts (Extended Data Table 2). Modelling was guided and validated by BS3 cross-linking data visualized with xVis^76^ and secondary structure predictions performed with Quick2D^77^ and PSIPRED^78, 79^. Amino acid residues connecting the domains of the two Rsc8 subunits could not be modelled. For clarification, they were placed into a single chain (chain L) clustered by proximity. The Sfh1 C-terminal finger helix was built into the density of map 7. A poly-alanine model was placed into density that could not be assigned to any RSC subunit (chain X). Bulky amino acid side chain density in the maps 4 – 8 enabled us to assign the sequence registers, however in some regions register shifts cannot be entirely excluded. The modelled RSC subunits Rsc4, Rsc58, Rsc6, Rsc8, Rsc9, Npl6, Htl1, Sfh1 and Sth1 (residues 48 – 293) together with the poly-alanine chain were applied to several rounds of real space refinement and geometry optimisation using PHENIX^67^, and flexible fitting with Namdinator^66^ against the combined map 8. MolProbity^80^ was used to flip and optimise Asn, Gln and His side chains. The C-terminal helix of Sfh1 was real space refined with PHENIX against map 7. The final structure displayed excellent stereochemistry as shown by MolProbity (Extended Data Table 3). Figures were created using PyMol^81^,UCSF Chimera^64^ and UCSF ChimeraX^82^. The angular distribution plots were generated using the AngularDistribution tool distributed with Warp^62^.

Sites of mutations found in human cancers were derived from the cBio cancer genomics portal (cBioPortal)^83, 84^ and mapped onto the RSC structure for residues that are identical in its human counterpart PBAF using MSAProbs^77, 85^. MSAPobs and Aline^86^ were used to map conservation between RSC and PBAF.

### Preparation of cross-linking samples for mass spectrometry

RSC-nucleosome complex was prepared as described above. The cross-linking reaction was performed with BS3 (bis(sulfosuccinimidyl(suberate))) cross-linker (Thermo Fischer Scientific) at a final concentration of 1 mM on ice for 30 min before quenching with Tris-HCl, pH 7.5, and ammonium bicarbonate at a final concentration of 100 mM and 20 mM, respectively. The cross-linked sample was applied to a 10% – 25% sucrose gradient as described above (no glutaraldehyde in the heavy solution) and protein containing fractions were pooled (∼800 µL, ∼50 µg complex) applied to in-solution digest. 150 µL of urea buffer (8 M urea, 50 mM NH_4_HCO_3_ pH 8) and 60 µL 0.1 M DTT (in 50 mM NH_4_HCO_3_ pH 8) were added to reduce the sample for 30 min at 37 °C, 300 rpm. The sample was alkylated with 60 µL 0.4 M iodoacetamide (in 50 mM NH_4_HCO_3_ pH 8) for 30 min at 37 °C, 300 rpm, in the dark. The reaction was quenched by addition of 60 µL 0.1 M DTT (in 50 mM NH_4_HCO_3_ pH 8). The sample was digested for 30 min at 37 °C with 0.5 µL Pierce Universal Nuclease (250 U/µl) in presence of 1 mM MgCl_2_. The final sample volume was adjusted to 1200 µL with 50 mM NH_4_HCO_3_ pH 8 resulting in a final urea concentration of 1 M. Trypsin digest was performed overnight at 37 °C with 2.5 µg trypsin (Promega, V5111). Tryptic peptides were desalted with C18 spin columns (Harvard Apparatus 74-4601), lyophilized and dissolved in 30% (v/v) acetonitrile, 0.1% (v/v) trifluoroacetic acid. The peptide mixture was separated on a Superdex Peptide 3.2/300 (GE Healthcare) column run at 50 µl/min with 30% (v/v) acetonitrile, 0.1% (v/v) trifluoroacetic acid. Cross-linked species are enriched by size exclusion chromatography based on their higher molecular weight compared to linear peptides. Therefore 50 µL fractions were collected from 1.0 mL post-injection. Fractions from 1.0 – 1.6 mL post-injection were dried in a speed-vac and dissolved in 5% (v/v) acetonitrile, 0.05% (v/v) trifluoroacetic acid and subjected to LC-MS/MS.

### LC-MS/MS analysis and cross-link identification

LC-MS/MS analyses were performed on a Q Exactive HF-X hybrid quadrupole-orbitrap mass spectrometer (Thermo Scientific) coupled to a Dionex Ultimate 3000 RSLCnano system. The peptide mixtures from in-solution digest were loaded on a Pepmap 300 C18 column (Thermo Fisher) at a flow rate of 10 µL/min in buffer A (0.1 % (v/v) formic acid) and washed for 3 min with buffer A. The sample was separated on an in-house packed C18 column (30 cm; ReproSil-Pur 120 Å, 1.9 µm, C18-AQ; inner diameter, 75 µm) at a flow rate of 300 nL/min. Sample separation was performed over 120 min using a buffer system consisting of 0.1 % (v/v) formic acid (buffer A) and 80 % (v/v) acetonitrile, 0.08 % (v/v) formic acid (buffer B). The main column was equilibrated with 5 % B, followed by sample application and a wash with 5 % B. Peptides were eluted by a linear gradient from 15 – 48 % B. The gradient was followed by a wash step at 95 % B and re-equilibration at 5 % B. Eluting peptides were analyzed in positive mode using a data-dependent top 30-acquisition methods. MS1 and MS2 resolution were set to 120,000 and 30,000 FWHM, respectively. Precursors selected for MS2 were fragmented using 30 % normalized, higher-energy collision-induced dissociation (HCD) fragmentation. Allowed charge states of selected precursors were +3 to +7. Further MS/MS parameters were set as follows: isolation width, 1.4 *m/z*; dynamic exclusion, 10 sec; max. injection time (MS1/MS2), 60 ms / 200 ms. The lock mass option (*m/z* 445.12002) was used for internal calibration. All measurements were performed in duplicates. The .raw files of all replicates were searched by the software pLink 2, version 2.3.1^87^ against a customized protein database containing the expressed proteins and protein-protein crosslinks were filtered with 1 % FDR. Cross-links appearing less than three times were excluded to increase confidence and plotted using xVis^76^ and xiNET^88^.

### Data availability statement

Coordinate file for RSC-nucleosome complex structure was deposited with the Protein Data Bank with accession codes XXXX. The cryo-EM density maps were deposited with the Electron Microscopy Data Base (EMDB) with accession codes EMD-XXXX etc.

## EXTENDED DATA FIGURE LEGENDS

**Extended Data Figure 1.**
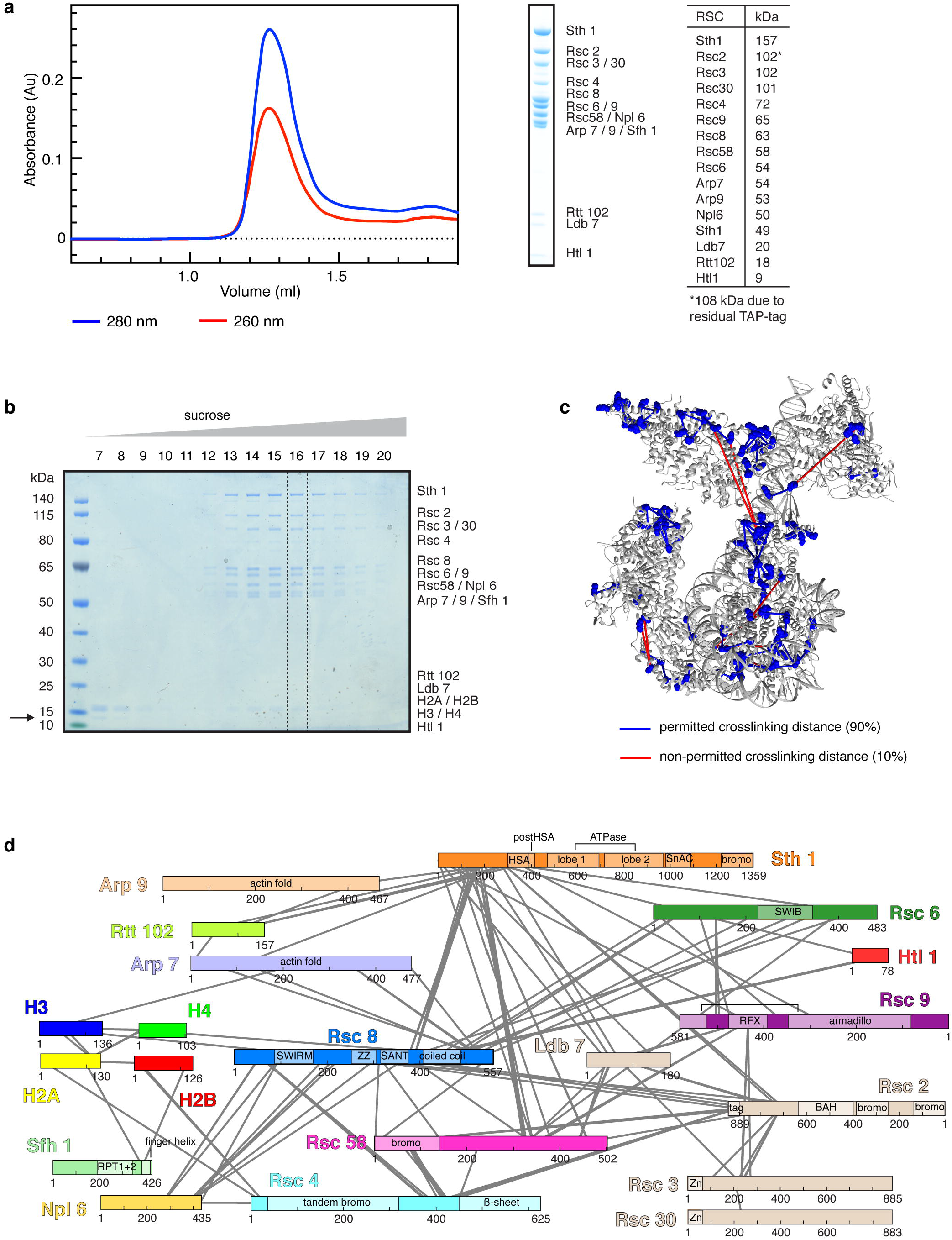
Preparation and characterization of RSC-nucleosome complex. Related to Figure 1. **a.** Preparation of endogenous Rsc2-containing isoform of the RSC complex from *S. cerevisiae*. Analysis of purified RSC by size-exclusion chromatography and SDS-PAGE showed high purity and homogeneity with stoichiometric subunits as assessable by Coomassie stain. Subunit identity was confirmed by mass spectrometry. The table shows the expected molecular weights of the RSC subunits. **b.** Assembly of the RSC-nucleosome complex. SDS-PAGE analysis of the fractions 7 – 20 of a 10 – 25% sucrose gradient ultracentrifugation. Complex formation was successful as demonstrated by the co-migration of histones with the RSC complex. The unbound over-stoichiometric nucleosomes only migrated to fraction 7 and 8 (black arrow). Fraction 16 in the presence of cross-linker was used for cryo-EM grid preparation (dashed box). **c.** Location of crosslinking sites mapped onto the structure. BS3 crosslinks that appeared at least in triplicates were mapped onto the RSC-nucleosome structure. Lysine residues involved in the crosslinking network are shown as blue spheres and crosslinked residues are connected with lines indicating permitted (blue) and non-permitted (red) crosslinking distances. 90% of the mapped crosslinks are within the permitted crosslinking distance which was set to 30 Å. The remaining 10% of non-permitted crosslinks likely reflect ambiguity caused by the presence of two identical Rsc8 subunits in the structure as well as flexibility of the complex in buffer or arise from technical errors. **d.** Crosslinking network between subunits of the RSC-nucleosome complex. Subunits are coloured as in Figure 1. Crosslinks with a score above 2.5 are shown. A comprehensive list of crosslinks can be found in the Supplementary Table S1.

**Extended Data Figure 2.**
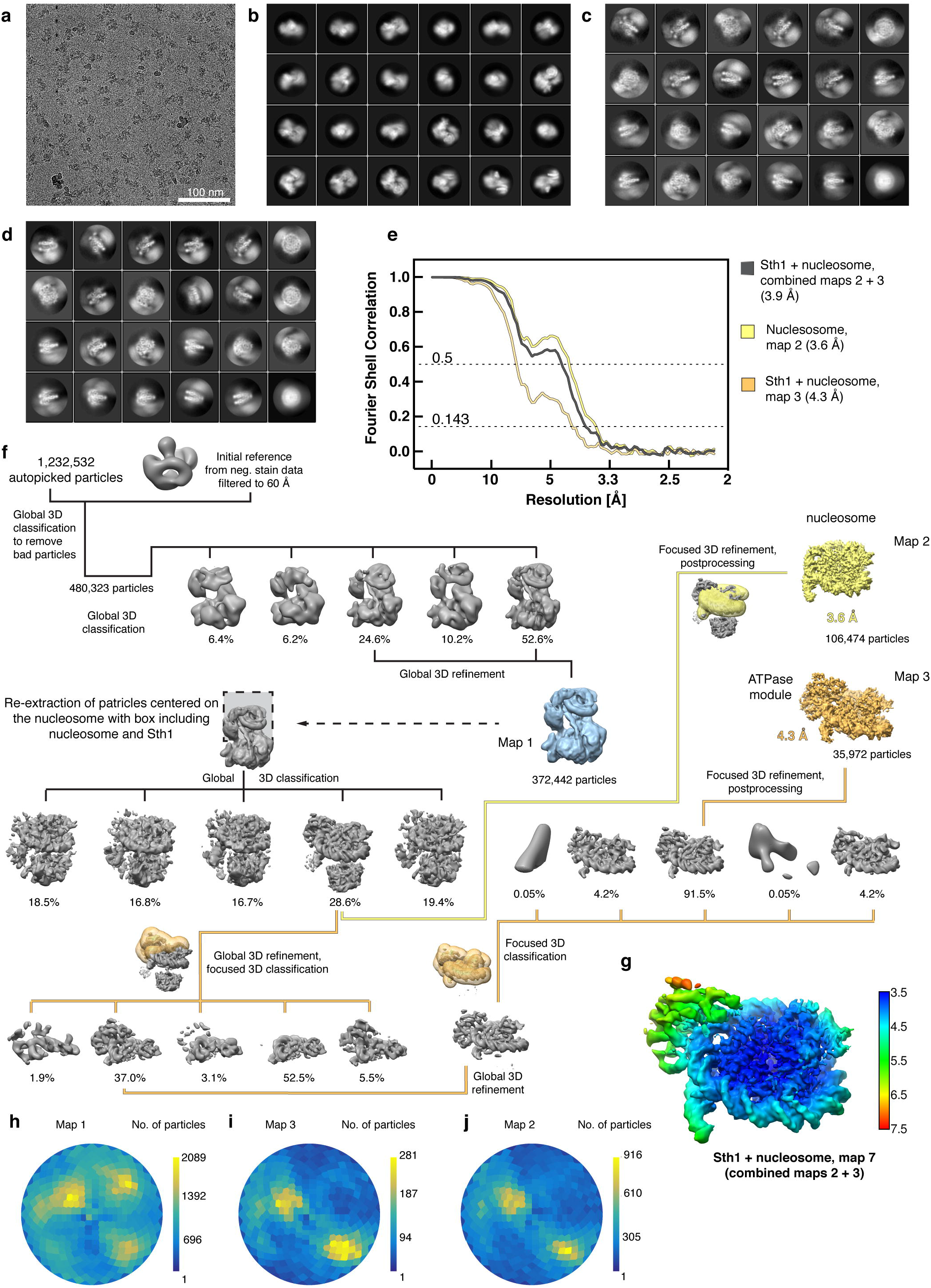
Cryo-EM analysis of the RSC-nucleosome complex. Related to Figures 1 – 6. **a**. Representative cryo-EM micrograph of the RSC-nucleosome complex shows homogeneously distributed individual particles. **b-d**. 2D class averages of the RSC-nucleosome (b), the Sth1-nucleosome subcomplex (c) and the nucleosome subcomplex (d). **e.** Fourier shell correlation plots reveal the overall resolutions of the cryo-EM reconstructions. **f.** Cryo-EM processing workflow for the reconstructions of the RSC-nucleosome, the Sth1-nucleosome subcomplex, and the nucleosome subcomplex. Particle distribution after 3D classifications is indicated below the corresponding map. The final maps are shown in colours. The masks used for focused classifications and refinements are colour coded corresponding to the final maps they were used for. Views are generally rotated by 180° with respect to Figure 1c, left. **g.** Local resolution estimation of the combined Sth1-nucleosome map as implemented in RELION^63^. We note that the resolution of the peripheral area with the Sth1 subunit is overestimated. **h-j**. Angular distribution plot for all particles contributing to the final reconstructions of the RSC-nucleosome (h), the Sth1-nucleosome (i) and the nucleosome complex (j).

**Extended Data Figure 3.**
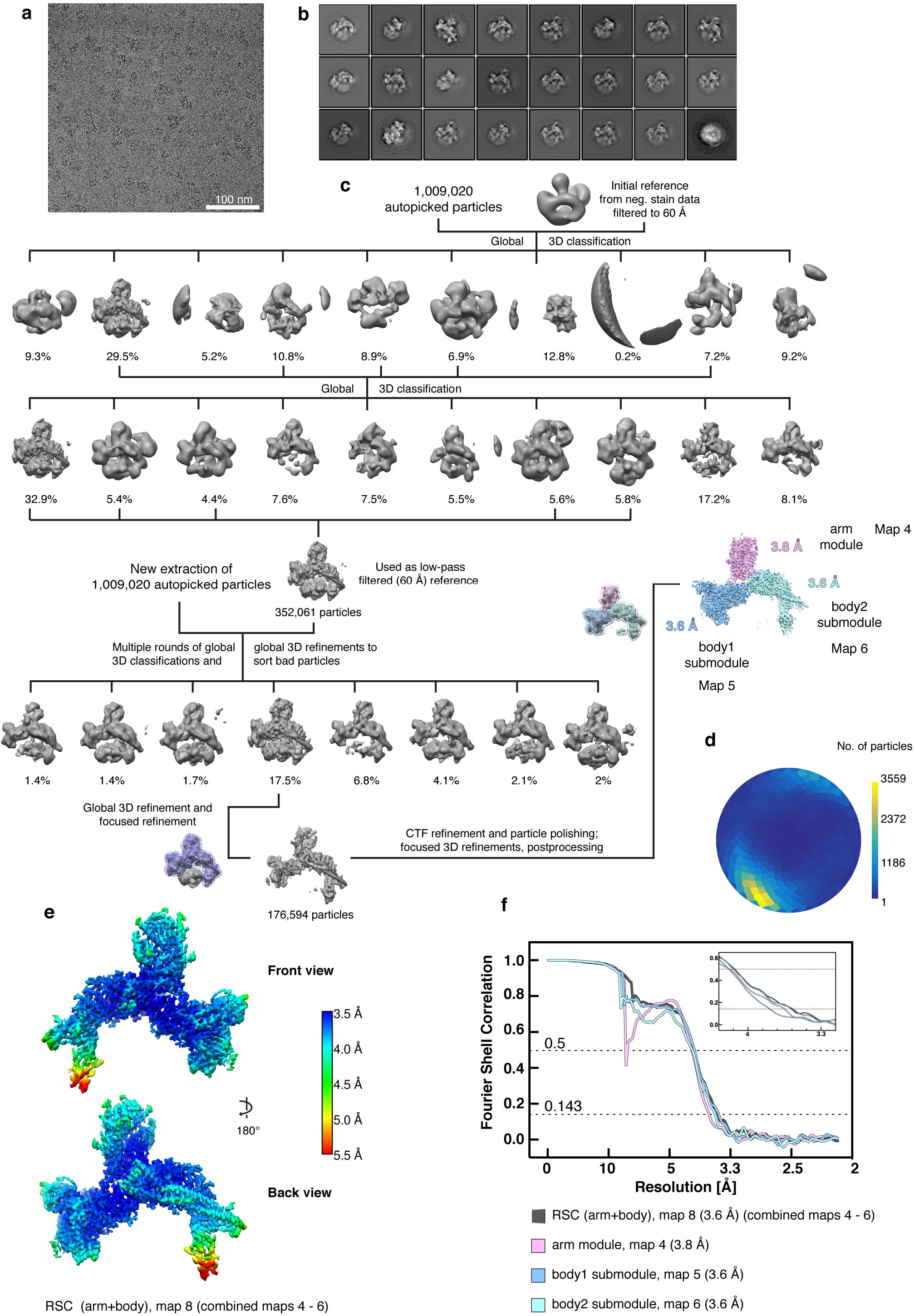
Cryo-EM analysis of the free RSC complex. Related to Figures 1 – 6. **a.** Representative cryo-EM micrograph of the free RSC complex shows homogeneously spaced individual particles. **b.** 2D class averages of the free RSC complex. **c.** Cryo-EM processing workflow for the reconstruction of the free RSC complex. Particle distribution after 3D classifications is indicated below the corresponding map. The final maps after focused 3D refinement and masks are depicted in colour. Views are generally rotated by 180° with respect to Figure 1c, right. **d.** Angular distribution plot for all particles contributing to the final reconstruction of the free RSC complex. **e.** Two views of the combined RSC core map coloured according to the local resolution as implemented in RELION^63^. **f.** Fourier shell correlation plots of the maps used for model building of the RSC core complex.

**Extended Data Figure 4.**
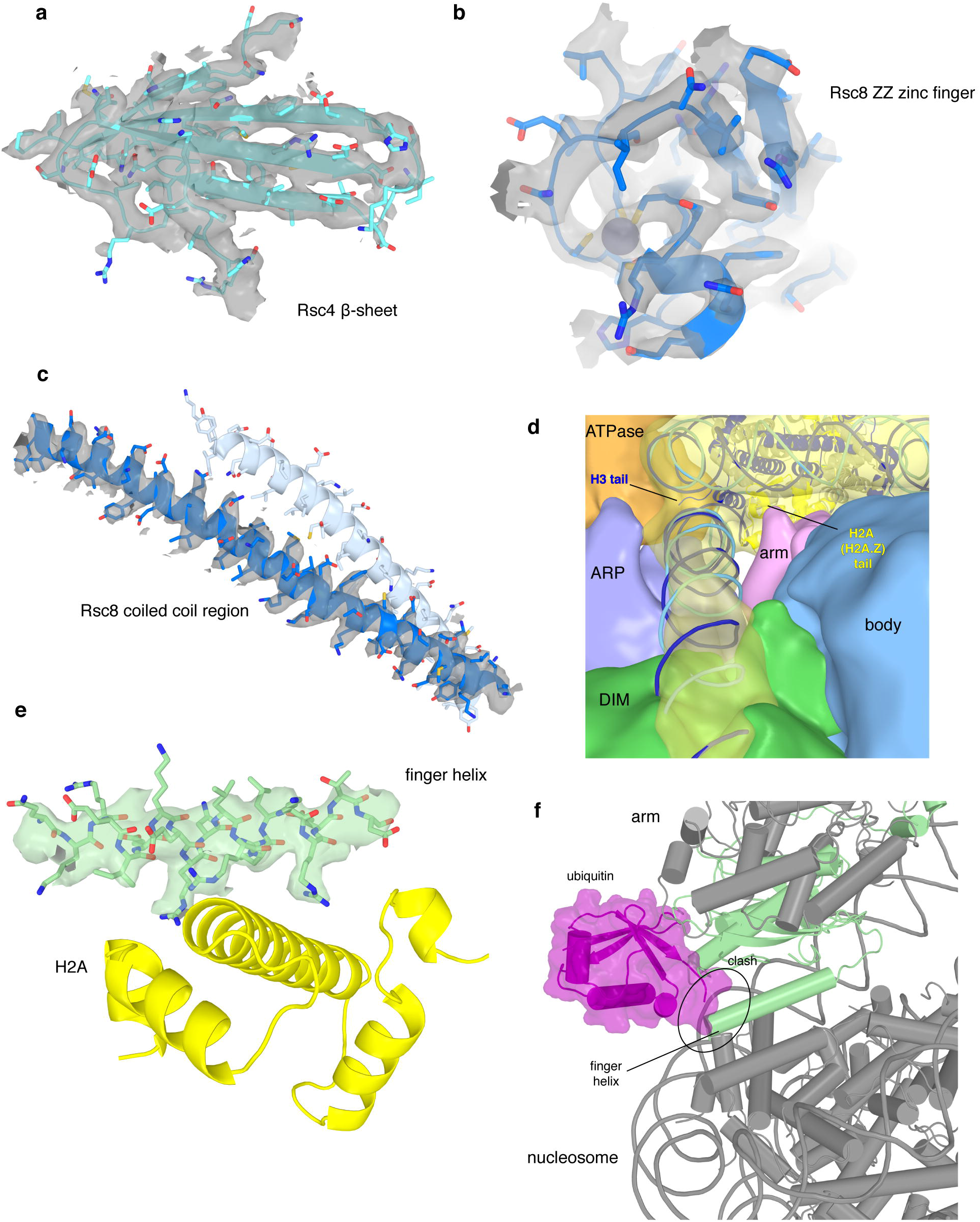
Cryo-EM densities for selected RSC regions. Related to Figures 1 – 4. **a-c**. Examples of map quality. Close-up of the Rsc4 β-sheet shows clear separation of individual strands (a). The high quality of the map for the ZZ zinc finger of Rsc8 allowed backbone tracing and placement of side chains as well as for the zinc ion (b). Coiled coil helices of the two Rsc8 subunits with density for one helix (c). **d.** View along the exit DNA in the direction of the nucleosome showing the low pass-filtered maps for the modules ATPase, ARP, DIM, arm, body, and the nucleosome. At the site where the H2A C-terminal tail protrudes from the nucleosome, low resolution density connecting the arm module and the nucleosome is visible. Density bridging form the ARP module to the exit DNA close to the H3 histone tail can be observed. **e.** Density representing the finger helix (green) at the acidic patch of the nucleosome (indicated by H2A in yellow). Side chain density is visible for conserved arginine residues. **f.** Interaction of RSC with the nucleosome is sterically impaired by H2B ubiquitylation at K120. The Sfh1 finger helix and the ubiquitin moiety (ubiquitylated nucleosome PDB code 6NOG)^89^ overlap after superposition of nucleosomes.

**Extended Data Figure 5.**
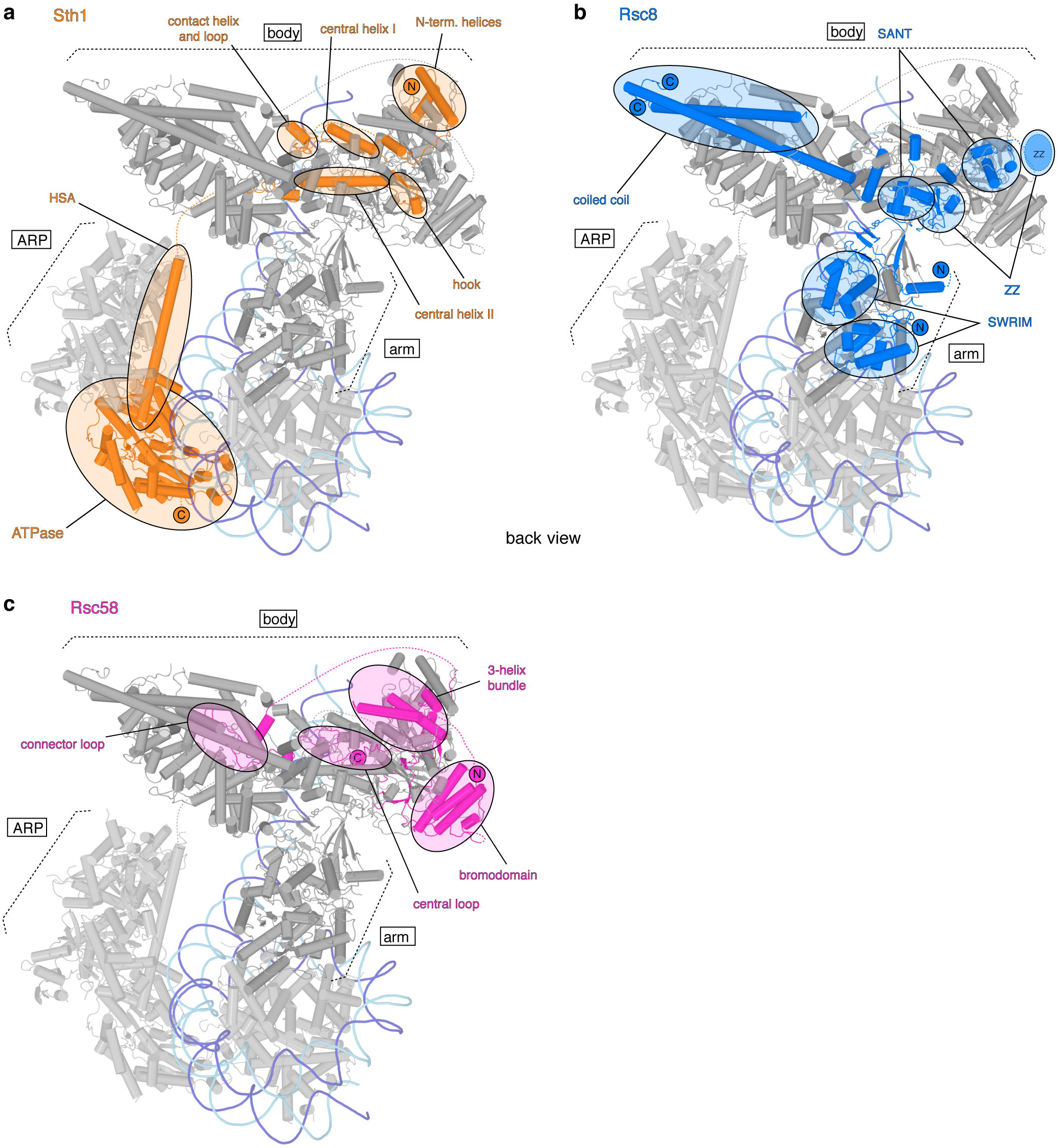
Course of polypeptide chains of architectural subunits Sth1, Rsc8 and Rsc58. Related to Figure 1. **a.** The Sth1 subunit of RSC starts with its N-terminus in the body module and tracks through it turning around with a contact helix and loop. Forming the central helix I, the hook and the central helix II it folds back and forth tightly interweaving the body module before it exits with its HSA region through the ARP module to build the ATPase module. **b.** Back view of the RSC remodeller with the domains of the two Rsc8 subunits highlighted in blue. Both Rsc8 start N-terminal with their SWIRM domains in the arm module where they support the two repeat domains of Sfh1 in a similar manner. They then follow distinct paths through the arm towards the body module where they contribute with both their SANT and ZZ zinc finger domains. Here the two domains of each subunit form different contacts with various interactions partners and whereas one ZZ zinc finger domain is tightly packed at the body and DNA-interaction module interface, the other seems to extend from the body, presumably as additional interaction surface. Both Rsc8 subunits unite again with their C-terminal long helices in a coiled coil fold in on the opposite side of the body module. **c.** Rsc58 N-terminal bromodomain attaches to the top of the body module. Then, Rsc58 follows an interwound path through the body module via the central and connector loop. It turns back docking to the body with a 3-helix bundle and stabilizing the module with its C-terminal end.

**Extended Data Figure 6.**
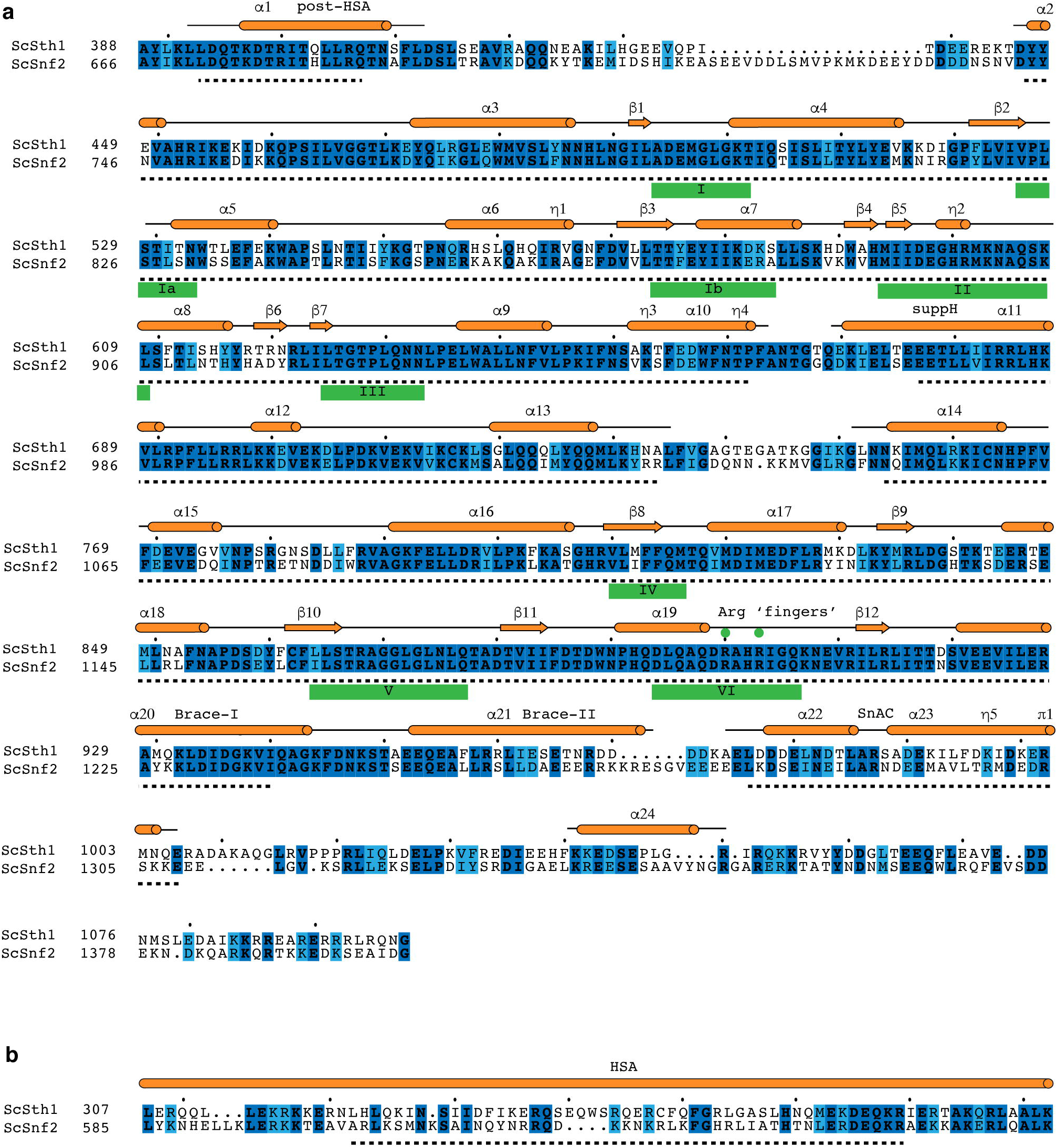
Sequence alignments for the Sth1 ATPase domain and HSA region. Related to Figures 1 – 6. **a.** Sequence alignment of the *S. cerevisiae* Sth1 ATPase domain to the homologous Snf2 ATPase domain of the same organism. Secondary structure elements are represented in red according to the cryo-EM structure of the Snf2 ATPase (PDB entry 5Z3U)^24^. Residues modelled in the Snf2 structure are topped by a back line with helical regions shown as cylinders and sheet regions as arrows. The Sth1 residues modelled in this work are indicated with a black dashed line below. ATPase motifs are underlined. Invariant residues are coloured in dark blue and conserved residues in light blue. The alignment was generated with MSAProbs^85^ within the MPI Bioinformatics Toolkit^77^ and visualized using ESPript^90^. **b.** Sequence alignment of the HSA regions from *S. cerevisiae* homologs Sth1 and Snf2. Illustration and generation of the alignment as in (a).

## EXTENDED DATA TABLES

**Extended Data Table 1.**
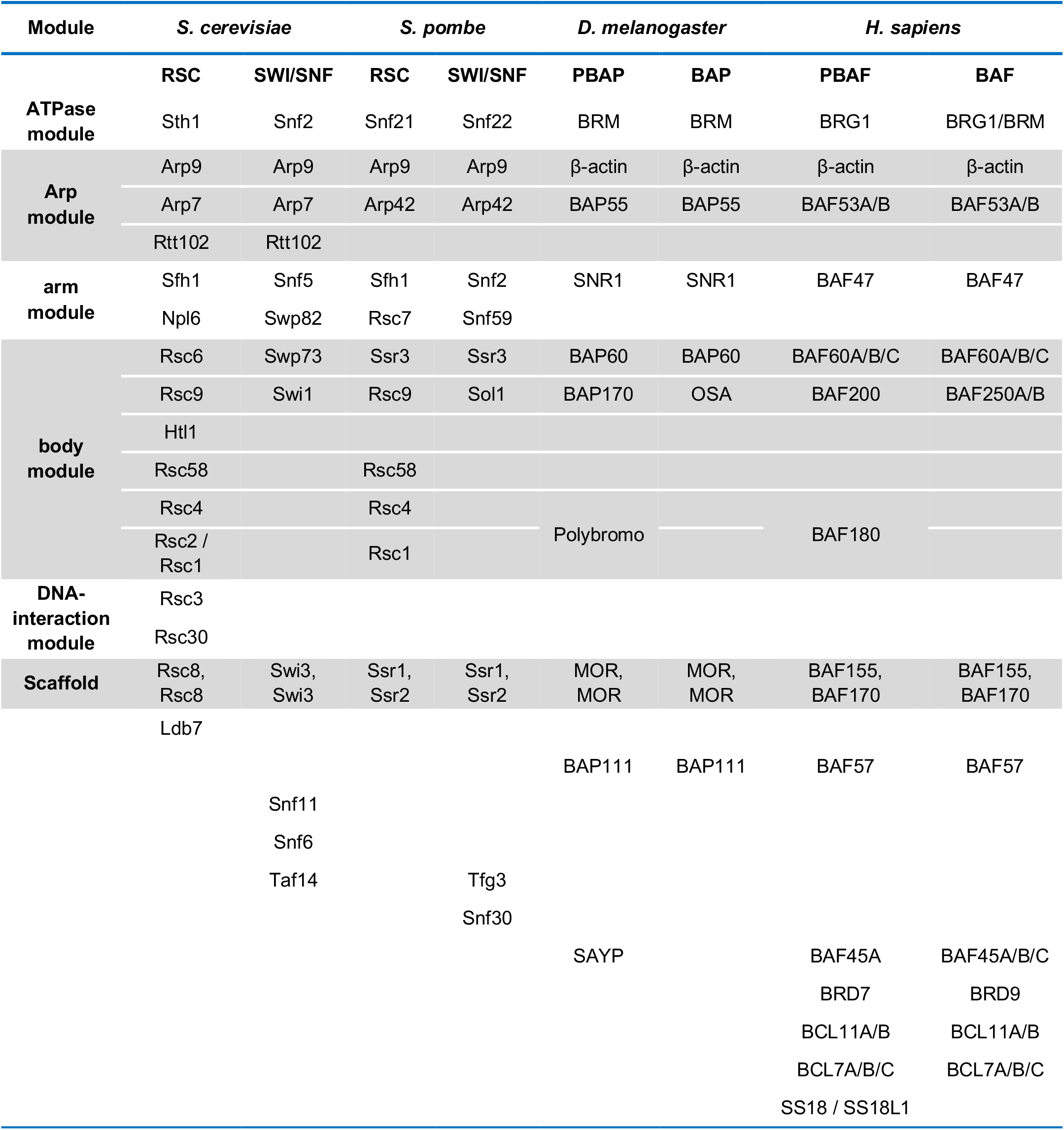
Subunit composition of RSC and related chromatin remodelling complexes. Assignment to the structural modules based on the *S. cerevisiae* structure of RSC presented in this work. Subunits occurring together in the complex are separated by comma, a slash indicates the use of one of the subunits. Subunits that could not be assigned to any module by homology are listed below. PBAF subunits contain 12 DNA-binding domains located in subunits BAF180 (HMG box)^91^, BAF200 (AT-rich domain, two C2H2 zinc fingers, RFX domain)^42^, BAF57 (HMG box)^92^ and BCL11A/B (six C2H2 zinc fingers)^93^.

**Extended Data Table 2.**
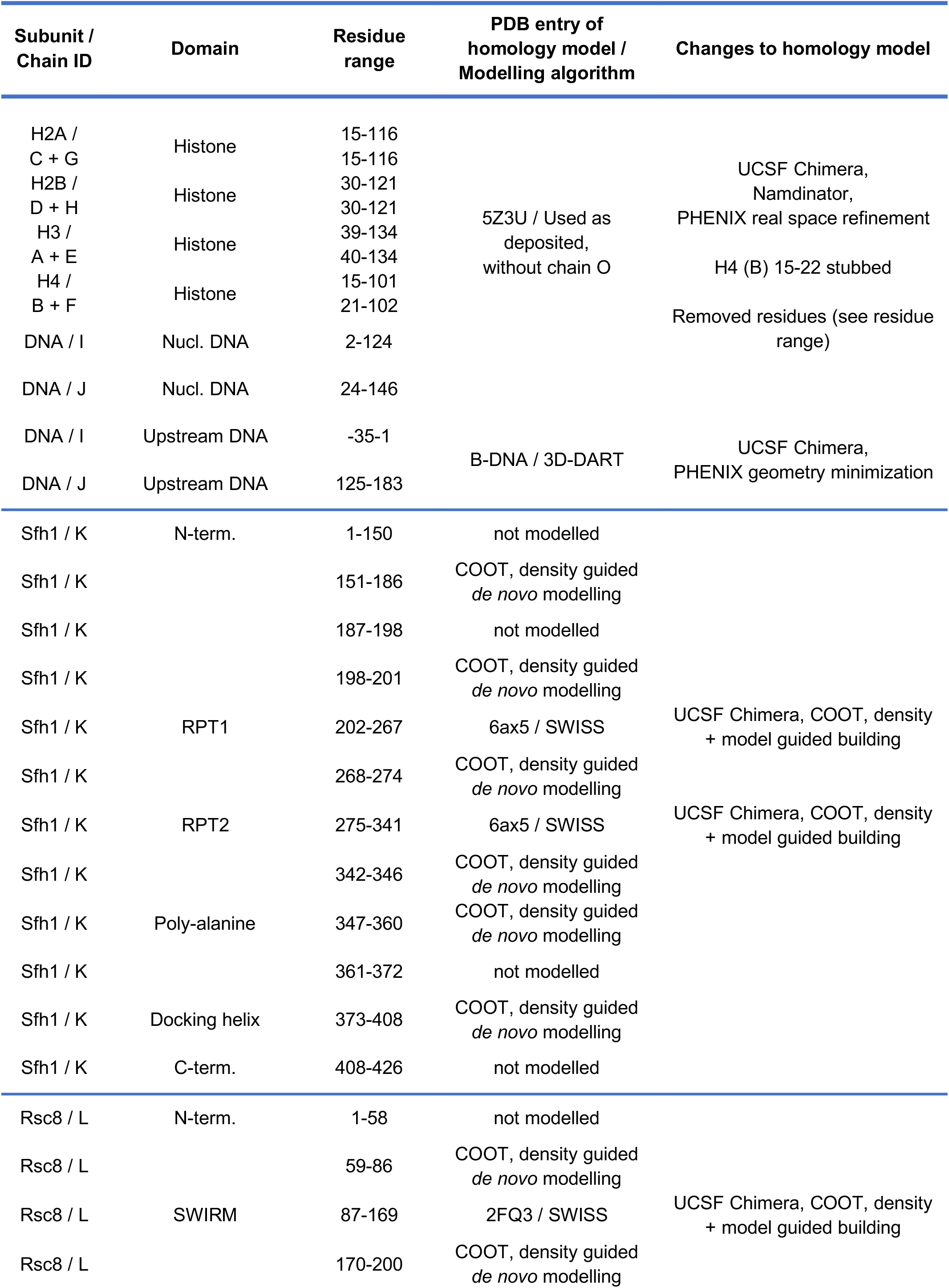
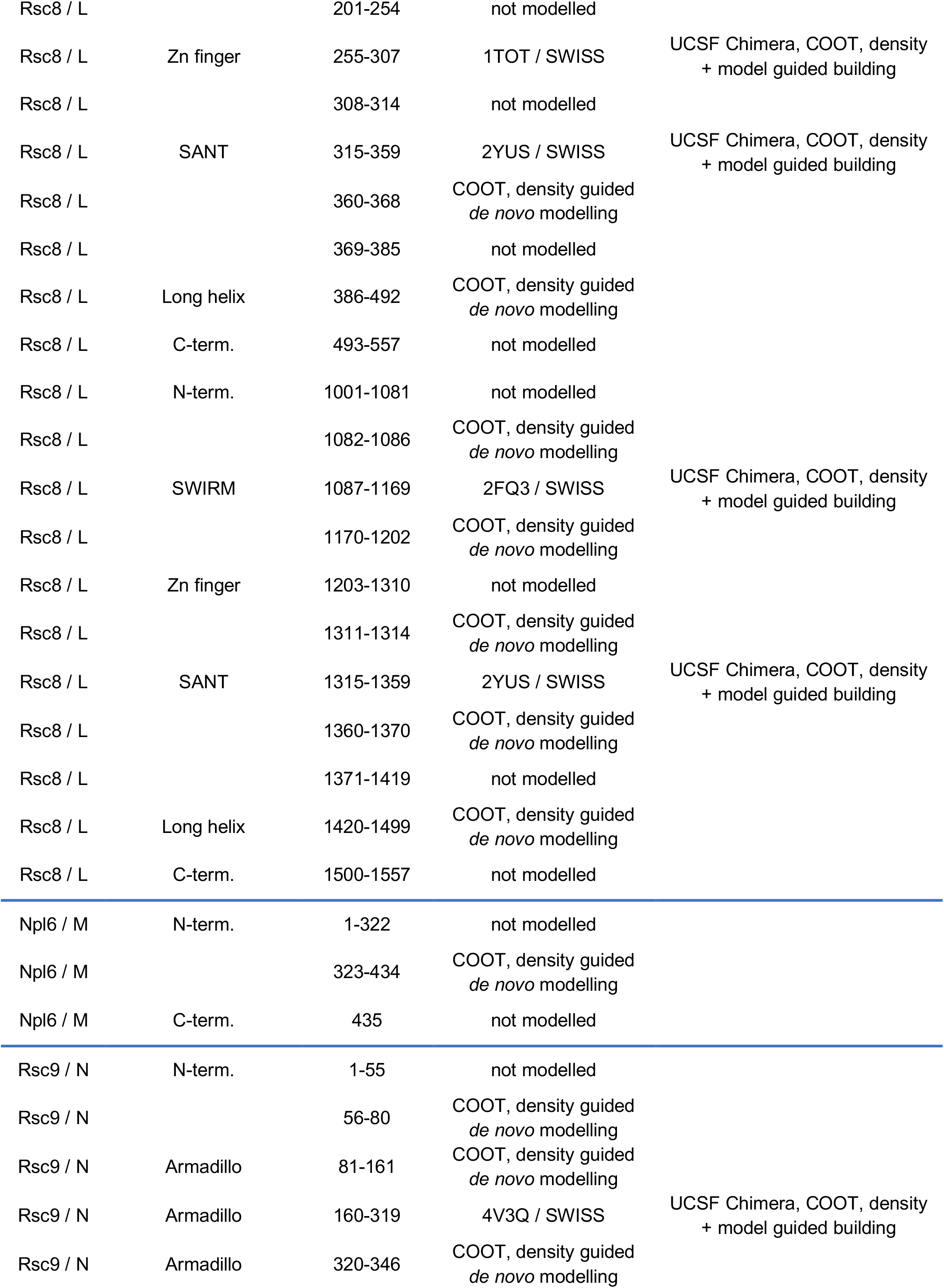
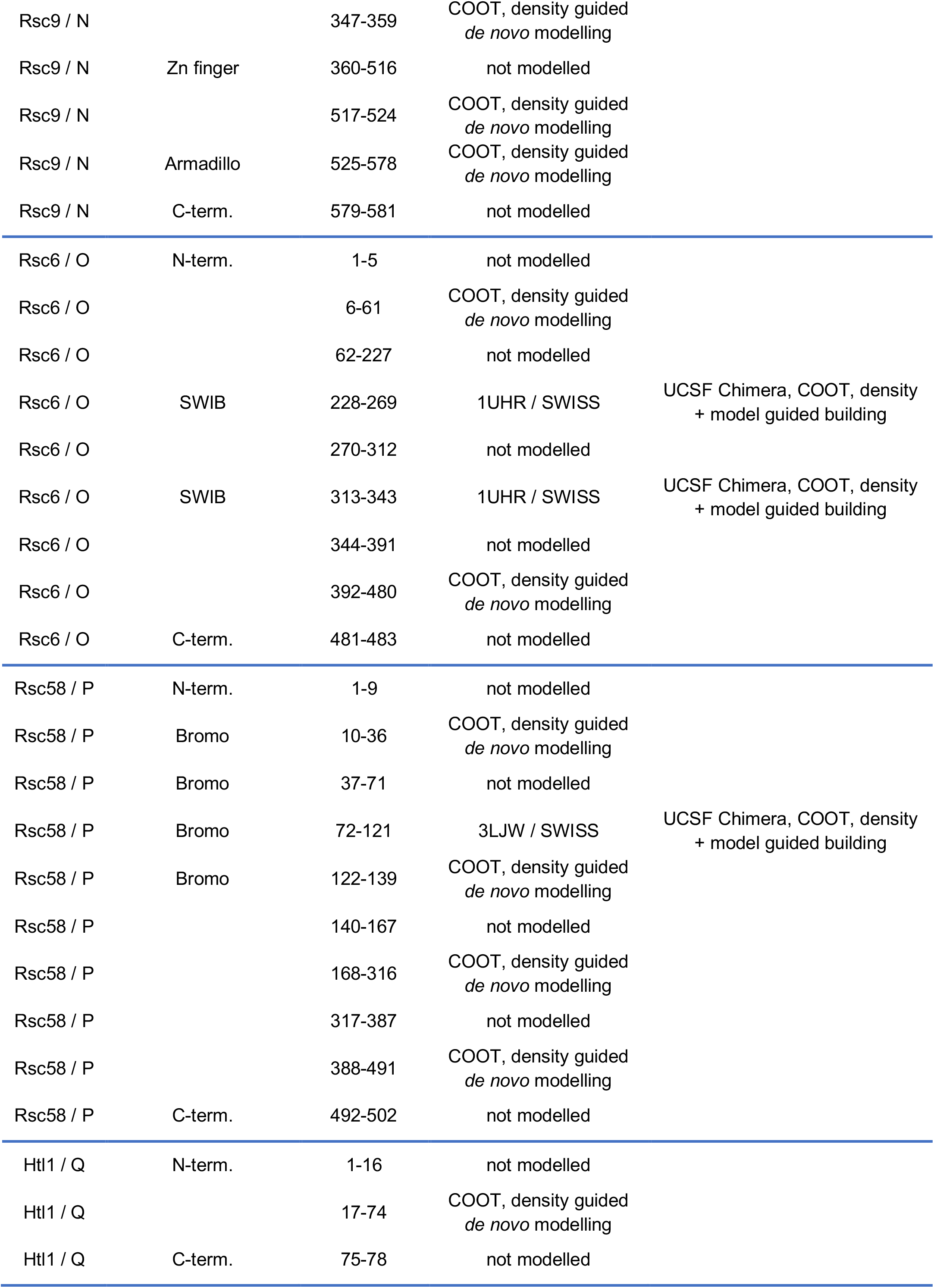
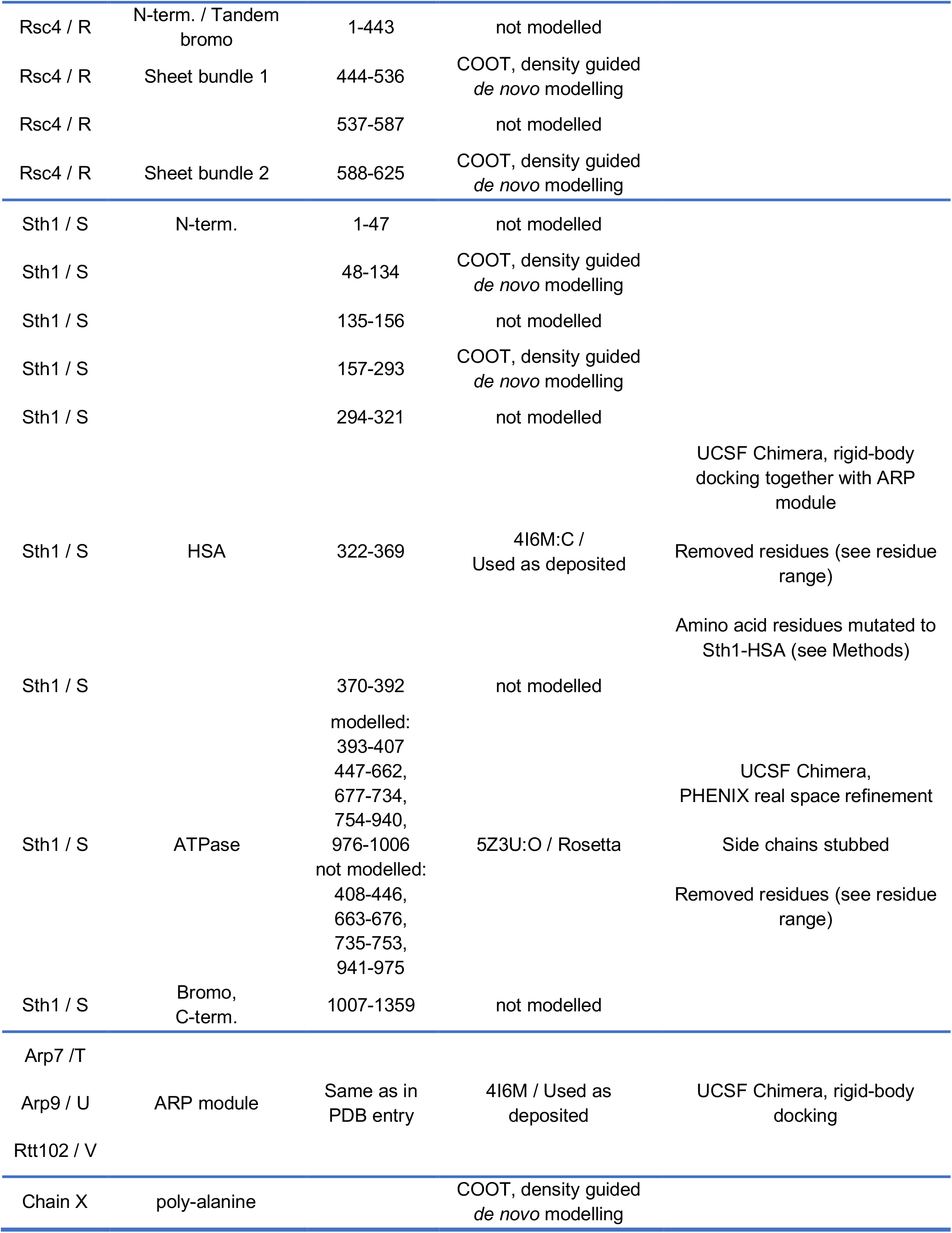
RSC subunit modelling. Modelling details for the RSC complex. Density that could not be assigned to a subunit was modelled with a poly alanine backbone in chain X. The domains of the two Rsc8 subunits are combined in one chain (L) spaced by 1000 amino acid offset.

**Extended Data Table 3.** Cryo-EM data collection, refinement, and validation statistics.

|  | Sth1-nucleosome structure<br>(map 7)<br>(EMDB-xxxx)<br>(PDB xxxx) | RSC core structure<br>(map 8)<br>(EMDB-xxxx)<br>(PDB xxxx) |
| --- | --- | --- |
| **Data collection and processing** |  |  |
| Magnification | 130,000 | 130,000 |
| Voltage (kV) | 300 | 300 |
| Electron exposure (e-/Å<sup>2</sup>) | 47.8 / 45.4* | 39.0<sup>†</sup> / 45.02<sup>†</sup> / 54.9<sup>‡</sup> |
| Defocus range (µm) | 0.4 – 2.9 | 0.8 – 4.8 |
| Pixel size (Å) | 1.05 | 1.05 |
| Symmetry imposed | C1 | C1 |
| Initial particle images (no.) | 1,232,532 | 1,009,020 |
| Final particle images (no.) | 106,474 / 35,972 | 176,594 |
| Map resolution (Å) | 3.9 | 3.6 |
| FSC threshold | 0.143 | 0.143 |
| Map resolution range (Å) | 3.5 – 7.5 | 3.5 – 5.5 |
| **Refinement** |  |  |
| Initial model used (PDB code) | 5Z3U | 3LJW, 1UHR, 2FQ3,<br>2YUS, 1TOT, 4V3Q,<br>6AX5 |
| Model resolution (Å) | 3.63 | 3.76 |
| FSC threshold | 0.5 | 0.5 |
| Map sharpening *B* factor (Å<sup>2</sup>) | -150 | -105.7 |
| Model composition |  |  |
| Non-hydrogen atoms | 13,655 | 20,545 |
| Protein / Nucleotide residues | 1,274 / 246 | 2,678 / - |
| Ligands | 0 | 1x Zn |
| *B* factors (Å<sup>2</sup>) |  |  |
| Protein | 106.53 | 53.27 |
| Ligand |  | 30.00 |
| Nucleotide | 119.35 |  |
| R.m.s. deviations |  |  |
| Bond lengths (Å) | 0.004 | 0.004 |
| Bond angles (°) | 0.748 | 0.931 |
| Validation |  |  |
| MolProbity score | 1.19 | 1.57 |
| Clashscore | 4.07 | 4.30 |
| Poor rotamers (%) | 0.00 | 0.14 |
| Ramachandran plot |  |  |
| Favored (%) | 98.07 | 94.84 |
| Allowed (%) | 1.93 | 5.16 |
| Disallowed (%) | 0.00 | 0.00 |

\* Dataset from two collections were combined
Datasets from three collections were combined, <sup>†</sup> not tilted, <sup>‡</sup>25° tilt

**Supplementary Table S1 | RSC-nucleosome cross-links**

List of intra- and inter-subunit lysine-lysine crosslinks as identified by LC-MS analyses and subsequent database search using pLink 2. The respective scores of cross-link identification are listed as well as the number of CSMs (cross-linked spectra matches).

**Supplementary Video 1 | Overview of RSC structure.**

The video shows the structure of RSC rotating around a vertical axis. It first depicts the low pass-filtered cryo-EM density, showing the five lobes of RSC and the nucleosome with exit DNA extending from it. It then shows the high-resolution cryo-EM maps for RSC modules, and finally the structural model as a ribbon representation with subunits in different colours (colour code as in Figure 1).

## REFERENCES

1. Lorch, Y. & Kornberg, R. D. Chromatin-remodeling for transcription. Q Rev Biophys 50, e5, doi:10.1017/S003358351700004X (2017).

2. Clapier, C. R., Iwasa, J., Cairns, B. R. & Peterson, C. L. Mechanisms of action and regulation of ATP-dependent chromatin-remodelling complexes. Nat Rev Mol Cell Biol 18, 407–422, doi:10.1038/nrm.2017.26 (2017).

3. Pulice, J. L. & Kadoch, C. Composition and Function of Mammalian SWI/SNF Chromatin Remodeling Complexes in Human Disease. Cold Spring Harb Symp Quant Biol 81, 53–60, doi:10.1101/sqb.2016.81.031021 (2016).

4. Henikoff, S. Mechanisms of Nucleosome Dynamics In Vivo. Cold Spring Harb Perspect Med 6, doi:10.1101/cshperspect.a026666 (2016).

5. Hirschhorn, J. N., Brown, S. A., Clark, C. D. & Winston, F. Evidence that SNF2/SWI2 and SNF5 activate transcription in yeast by altering chromatin structure. Genes Dev 6, 2288–2298, doi:10.1101/gad.6.12a.2288 (1992).

6. Peterson, C. L. & Herskowitz, I. Characterization of the yeast SWI1, SWI2, and SWI3 genes, which encode a global activator of transcription. Cell 68, 573-583, doi:10.1016/0092-8674(92)90192-f (1992).

7. Cairns, B. R. et al. RSC, an essential, abundant chromatin-remodeling complex. Cell 87, 1249–1260, doi:10.1016/s0092-8674(00)81820-6 (1996).

8. Saha, A., Wittmeyer, J. & Cairns, B. R. Chromatin remodeling by RSC involves ATP-dependent DNA translocation. Genes Dev 16, 2120–2134, doi:10.1101/gad.995002 (2002).

9. Clapier, C. R. et al. Regulation of DNA Translocation Efficiency within the Chromatin Remodeler RSC/Sth1 Potentiates Nucleosome Sliding and Ejection. Mol Cell 62, 453–461, doi:10.1016/j.molcel.2016.03.032 (2016).

10. Szerlong, H. et al. The HSA domain binds nuclear actin-related proteins to regulate chromatin-remodeling ATPases. Nat Struct Mol Biol 15, 469–476, doi:10.1038/nsmb.1403 (2008).

11. Parnell, T. J., Huff, J. T. & Cairns, B. R. RSC regulates nucleosome positioning at Pol II genes and density at Pol III genes. EMBO J 27, 100–110, doi:10.1038/sj.emboj.7601946 (2008).

12. Krietenstein, N. et al. Genomic Nucleosome Organization Reconstituted with Pure Proteins. Cell 167, 709–721 e712, doi:10.1016/j.cell.2016.09.045 (2016).

13. Ng, H. H., Robert, F., Young, R. A. & Struhl, K. Genome-wide location and regulated recruitment of the RSC nucleosome-remodeling complex. Genes Dev 16, 806–819, doi:10.1101/gad.978902 (2002).

14. Klein-Brill, A., Joseph-Strauss, D., Appleboim, A. & Friedman, N. Dynamics of Chromatin and Transcription during Transient Depletion of the RSC Chromatin Remodeling Complex. Cell Rep 26, 279–292 e275, doi:10.1016/j.celrep.2018.12.020 (2019).

15. Brahma, S. & Henikoff, S. RSC-Associated Subnucleosomes Define MNase-Sensitive Promoters in Yeast. Mol Cell 73, 238–249 e233, doi:10.1016/j.molcel.2018.10.046 (2019).

16. Kubik, S. et al. Nucleosome Stability Distinguishes Two Different Promoter Types at All Protein-Coding Genes in Yeast. Mol Cell 60, 422–434, doi:10.1016/j.molcel.2015.10.002 (2015).

17. Ramachandran, S., Zentner, G. E. & Henikoff, S. Asymmetric nucleosomes flank promoters in the budding yeast genome. Genome Res 25, 381–390, doi:10.1101/gr.182618.114 (2015).

18. Badis, G. et al. A library of yeast transcription factor motifs reveals a widespread function for Rsc3 in targeting nucleosome exclusion at promoters. Mol Cell 32, 878–887, doi:10.1016/j.molcel.2008.11.020 (2008).

19. Lorch, Y., Maier-Davis, B. & Kornberg, R. D. Role of DNA sequence in chromatin remodeling and the formation of nucleosome-free regions. Genes Dev 28, 2492–2497, doi:10.1101/gad.250704.114 (2014).

20. Kubik, S. et al. Sequence-Directed Action of RSC Remodeler and General Regulatory Factors Modulates +1 Nucleosome Position to Facilitate Transcription. Mol Cell 71, 89–102 e105, doi:10.1016/j.molcel.2018.05.030 (2018).

21. Asturias, F. J., Chung, W. H., Kornberg, R. D. & Lorch, Y. Structural analysis of the RSC chromatin-remodeling complex. Proc Natl Acad Sci U S A 99, 13477–13480, doi:10.1073/pnas.162504299 (2002).

22. Chaban, Y. et al. Structure of a RSC-nucleosome complex and insights into chromatin remodeling. Nat Struct Mol Biol 15, 1272–1277, doi:10.1038/nsmb.1524 (2008).

23. Leschziner, A. E. et al. Conformational flexibility in the chromatin remodeler RSC observed by electron microscopy and the orthogonal tilt reconstruction method. Proc Natl Acad Sci U S A 104, 4913–4918, doi:10.1073/pnas.0700706104 (2007).

24. Li, M. et al. Mechanism of DNA translocation underlying chromatin remodelling by Snf2. Nature 567, 409–413, doi:10.1038/s41586-019-1029-2 (2019).

25. Schubert, H. L. et al. Structure of an actin-related subcomplex of the SWI/SNF chromatin remodeler. Proc Natl Acad Sci U S A 110, 3345–3350, doi:10.1073/pnas.1215379110 (2013).

26. Kasten, M. et al. Tandem bromodomains in the chromatin remodeler RSC recognize acetylated histone H3 Lys14. EMBO J 23, 1348–1359, doi:10.1038/sj.emboj.7600143 (2004).

27. Treich, I., Ho, L. & Carlson, M. Direct interaction between Rsc6 and Rsc8/Swh3, two proteins that are conserved in SWI/SNF-related complexes. Nucleic Acids Res 26, 3739–3745, doi:10.1093/nar/26.16.3739 (1998).

28. Taneda, T. & Kikuchi, A. Genetic analysis of RSC58, which encodes a component of a yeast chromatin remodeling complex, and interacts with the transcription factor Swi6. Mol Genet Genomics 271, 479–489, doi:10.1007/s00438-004-0999-3 (2004).

29. Chambers, A. L., Pearl, L. H., Oliver, A. W. & Downs, J. A. The BAH domain of Rsc2 is a histone H3 binding domain. Nucleic Acids Res 41, 9168–9182, doi:10.1093/nar/gkt662 (2013).

30. VanDemark, A. P. et al. Autoregulation of the rsc4 tandem bromodomain by gcn5 acetylation. Mol Cell 27, 817–828, doi:10.1016/j.molcel.2007.08.018 (2007).

31. Saha, A., Wittmeyer, J. & Cairns, B. R. Chromatin remodeling through directional DNA translocation from an internal nucleosomal site. Nat Struct Mol Biol 12, 747–755, doi:10.1038/nsmb973 (2005).

32. Cairns, B. R., Erdjument-Bromage, H., Tempst, P., Winston, F. & Kornberg, R. D. Two actin-related proteins are shared functional components of the chromatin-remodeling complexes RSC and SWI/SNF. Mol Cell 2, 639–651 (1998).

33. Sen, P. et al. The SnAC domain of SWI/SNF is a histone anchor required for remodeling. Mol Cell Biol 33, 360–370, doi:10.1128/MCB.00922-12 (2013).

34. Lorch, Y., Maier-Davis, B. & Kornberg, R. D. Histone Acetylation Inhibits RSC and Stabilizes the +1 Nucleosome. Mol Cell 72, 594–600 e592, doi:10.1016/j.molcel.2018.09.030 (2018).

35. Cao, Y., Cairns, B. R., Kornberg, R. D. & Laurent, B. C. Sfh1p, a component of a novel chromatin-remodeling complex, is required for cell cycle progression. Mol Cell Biol 17, 3323–3334, doi:10.1128/mcb.17.6.3323 (1997).

36. Sen, P., Ghosh, S., Pugh, B. F. & Bartholomew, B. A new, highly conserved domain in Swi2/Snf2 is required for SWI/SNF remodeling. Nucleic Acids Res 39, 9155–9166, doi:10.1093/nar/gkr622 (2011).

37. Cakiroglu, A. et al. Genome-wide reconstitution of chromatin transactions reveals that RSC preferentially disrupts H2AZ-containing nucleosomes. Genome Res 29, 988–998, doi:10.1101/gr.243139.118 (2019).

38. Suto, R. K., Clarkson, M. J., Tremethick, D. J. & Luger, K. Crystal structure of a nucleosome core particle containing the variant histone H2A.Z. Nat Struct Biol 7, 1121–1124, doi:10.1038/81971 (2000).

39. Materne, P. et al. Histone H2B ubiquitylation represses gametogenesis by opposing RSC-dependent chromatin remodeling at the ste11 master regulator locus. Elife 5, doi:10.7554/eLife.13500 (2016).

40. Angus-Hill, M. L. et al. A Rsc3/Rsc30 zinc cluster dimer reveals novel roles for the chromatin remodeler RSC in gene expression and cell cycle control. Mol Cell 7, 741–751 (2001).

41. Brogaard, K., Xi, L., Wang, J. P. & Widom, J. A map of nucleosome positions in yeast at base-pair resolution. Nature 486, 496–501, doi:10.1038/nature11142 (2012).

42. Yan, Z. et al. PBAF chromatin-remodeling complex requires a novel specificity subunit, BAF200, to regulate expression of selective interferon-responsive genes. Genes Dev 19, 1662-1667, doi:10.1101/gad.1323805 (2005).

43. Waterhouse, A. et al. SWISS-MODEL: homology modelling of protein structures and complexes. Nucleic Acids Res 46, W296–W303, doi:10.1093/nar/gky427 (2018).

44. Xue, Y. et al. The human SWI/SNF-B chromatin-remodeling complex is related to yeast rsc and localizes at kinetochores of mitotic chromosomes. Proc Natl Acad Sci U S A 97, 13015–13020, doi:10.1073/pnas.240208597 (2000).

45. Sandhya, S., Maulik, A., Giri, M. & Singh, M. Domain architecture of BAF250a reveals the ARID and ARM-repeat domains with implication in function and assembly of the BAF remodeling complex. PLoS One 13, e0205267, doi:10.1371/journal.pone.0205267 (2018).

46. Yan, L., Wu, H., Li, X., Gao, N. & Chen, Z. Structures of the ISWI-nucleosome complex reveal a conserved mechanism of chromatin remodeling. Nat Struct Mol Biol 26, 258–266, doi:10.1038/s41594-019-0199-9 (2019).

47. Farnung, L., Vos, S. M., Wigge, C. & Cramer, P. Nucleosome-Chd1 structure and implications for chromatin remodelling. Nature 550, 539–542, doi:10.1038/nature24046 (2017).

48. Sundaramoorthy, R. et al. Structure of the chromatin remodelling enzyme Chd1 bound to a ubiquitinylated nucleosome. Elife 7, doi:10.7554/eLife.35720 (2018).

49. Willhoft, O. et al. Structure and dynamics of the yeast SWR1-nucleosome complex. Science 362, doi:10.1126/science.aat7716 (2018).

50. Ayala, R. et al. Structure and regulation of the human INO80-nucleosome complex. Nature 556, 391–395, doi:10.1038/s41586-018-0021-6 (2018).

51. Eustermann, S. et al. Structural basis for ATP-dependent chromatin remodelling by the INO80 complex. Nature 556, 386–390, doi:10.1038/s41586-018-0029-y (2018).

52. Knoll, K. R. et al. The nuclear actin-containing Arp8 module is a linker DNA sensor driving INO80 chromatin remodeling. Nat Struct Mol Biol 25, 823–832, doi:10.1038/s41594-018-0115-8 (2018).

## METHODS AND EXTENDED DATA REFERENCES

53. Cairns, B. R. et al. Two functionally distinct forms of the RSC nucleosome-remodeling complex, containing essential AT hook, BAH, and bromodomains. Mol Cell 4, 715–723 (1999).

54. Rigaut, G. et al. A generic protein purification method for protein complex characterization and proteome exploration. Nat Biotechnol 17, 1030–1032, doi:10.1038/13732 (1999).

55. Lorch, Y. & Kornberg, R. D. Isolation and assay of the RSC chromatin-remodeling complex from Saccharomyces cerevisiae. Methods Enzymol 377, 316–322, doi:10.1016/S0076-6879(03)77019-0 (2004).

56. Luger, K., Rechsteiner, T. J. & Richmond, T. J. Expression and purification of recombinant histones and nucleosome reconstitution. Methods Mol Biol 119, 1–16, doi:10.1385/1-59259-681-9:1 (1999).

57. Dyer, P. N. et al. in Methods in Enzymology Vol. 375 23–44 (Academic Press, 2003).

58. Maskell, D. P. et al. Structural basis for retroviral integration into nucleosomes. Nature 523, 366–369, doi:10.1038/nature14495 (2015).

59. Lowary, P. T. & Widom, J. New DNA sequence rules for high affinity binding to histone octamer and sequence-directed nucleosome positioning. J Mol Biol 276, 19–42, doi:10.1006/jmbi.1997.1494 (1998).

60. Kastner, B. et al. GraFix: sample preparation for single-particle electron cryomicroscopy. Nat Methods 5, 53–55, doi:10.1038/nmeth1139 (2008).

61. Stark, H. GraFix: stabilization of fragile macromolecular complexes for single particle cryo-EM. Methods Enzymol 481, 109–126, doi:10.1016/S0076-6879(10)81005-5 (2010).

62. Tegunov, D. & Cramer, P. Real-time cryo-EM data pre-processing with Warp. bioRxiv (2018).

63. Zivanov, J. et al. New tools for automated high-resolution cryo-EM structure determination in RELION-3. Elife 7, doi:10.7554/eLife.42166 (2018).

64. Pettersen, E. F. et al. UCSF Chimera--a visualization system for exploratory research and analysis. J Comput Chem 25, 1605–1612, doi:10.1002/jcc.20084 (2004).

65. Emsley, P., Lohkamp, B., Scott, W. G. & Cowtan, K. Features and development of Coot. Acta Crystallogr D Biol Crystallogr 66, 486–501, doi:10.1107/S0907444910007493 (2010).

66. Kidmose, R. T. et al. Namdinator - automatic molecular dynamics flexible fitting of structural models into cryo-EM and crystallography experimental maps. IUCrJ 6, 526–531, doi:10.1107/S2052252519007619 (2019).

67. Adams, P. D. et al. PHENIX: a comprehensive Python-based system for macromolecular structure solution. Acta Crystallogr D Biol Crystallogr 66, 213–221, doi:10.1107/S0907444909052925 (2010).

68. Song, Y. et al. High-resolution comparative modeling with RosettaCM. Structure 21, 1735–1742, doi:10.1016/j.str.2013.08.005 (2013).

69. Raman, S. et al. Structure prediction for CASP8 with all-atom refinement using Rosetta. Proteins 77 Suppl 9, 89–99, doi:10.1002/prot.22540 (2009).

70. van Dijk, M. & Bonvin, A. M. 3D-DART: a DNA structure modelling server. Nucleic Acids Res 37, W235–239, doi:10.1093/nar/gkp287 (2009).

71. Bienert, S. et al. The SWISS-MODEL Repository-new features and functionality. Nucleic Acids Res 45, D313–D319, doi:10.1093/nar/gkw1132 (2017).

72. Charlop-Powers, Z., Zeng, L., Zhang, Q. & Zhou, M. M. Structural insights into selective histone H3 recognition by the human Polybromo bromodomain 2. Cell Res 20, 529–538, doi:10.1038/cr.2010.43 (2010).

73. Da, G. et al. Structure and function of the SWIRM domain, a conserved protein module found in chromatin regulatory complexes. Proc Natl Acad Sci U S A 103, 2057–2062, doi:10.1073/pnas.0510949103 (2006).

74. Legge, G. B. et al. ZZ domain of CBP: an unusual zinc finger fold in a protein interaction module. J Mol Biol 343, 1081–1093, doi:10.1016/j.jmb.2004.08.087 (2004).

75. Reichen, C. et al. Structures of designed armadillo-repeat proteins show propagation of inter-repeat interface effects. Acta Crystallogr D Struct Biol 72, 168–175, doi:10.1107/S2059798315023116 (2016).

76. Grimm, M., Zimniak, T., Kahraman, A. & Herzog, F. xVis: a web server for the schematic visualization and interpretation of crosslink-derived spatial restraints. Nucleic Acids Res 43, W362–369, doi:10.1093/nar/gkv463 (2015).

77. Zimmermann, L. et al. A Completely Reimplemented MPI Bioinformatics Toolkit with a New HHpred Server at its Core. J Mol Biol 430, 2237–2243, doi:10.1016/j.jmb.2017.12.007 (2018).

78. Buchan, D. W. A. & Jones, D. T. The PSIPRED Protein Analysis Workbench: 20 years on. Nucleic Acids Res 47, W402–W407, doi:10.1093/nar/gkz297 (2019).

79. Jones, D. T. Protein secondary structure prediction based on position-specific scoring matrices. J Mol Biol 292, 195–202, doi:10.1006/jmbi.1999.3091 (1999).

80. Williams, C. J. et al. MolProbity: More and better reference data for improved all-atom structure validation. Protein Sci 27, 293–315, doi:10.1002/pro.3330 (2018).

81. Schrodinger, LLC. The PyMOL Molecular Graphics System, Version 1.8 (2015).

82. Goddard, T. D. et al. UCSF ChimeraX: Meeting modern challenges in visualization and analysis. Protein Sci 27, 14–25, doi:10.1002/pro.3235 (2018).

83. Cerami, E. et al. The cBio cancer genomics portal: an open platform for exploring multidimensional cancer genomics data. Cancer Discov 2, 401–404, doi:10.1158/2159-8290.CD-12-0095 (2012).

84. Gao, J. et al. Integrative analysis of complex cancer genomics and clinical profiles using the cBioPortal. Sci Signal 6, pl1, doi:10.1126/scisignal.2004088 (2013).

85. Liu, Y., Schmidt, B. & Maskell, D. L. MSAProbs: multiple sequence alignment based on pair hidden Markov models and partition function posterior probabilities. Bioinformatics 26, 1958–1964, doi:10.1093/bioinformatics/btq338 (2010).

86. Bond, C. S. & Schuttelkopf, A. W. ALINE: a WYSIWYG protein-sequence alignment editor for publication-quality alignments. Acta Crystallogr D Biol Crystallogr 65, 510–512, doi:10.1107/S0907444909007835 (2009).

87. Yang, B. et al. Identification of cross-linked peptides from complex samples. Nat Methods 9, 904–906, doi:10.1038/nmeth.2099 (2012).

88. Combe, C. W., Fischer, L. & Rappsilber, J. xiNET: cross-link network maps with residue resolution. Mol Cell Proteomics 14, 1137–1147, doi:10.1074/mcp.O114.042259 (2015).

89. Worden, E. J., Hoffmann, N. A., Hicks, C. W. & Wolberger, C. Mechanism of Cross-talk between H2B Ubiquitination and H3 Methylation by Dot1L. Cell 176, 1490–1501 e1412, doi:10.1016/j.cell.2019.02.002 (2019).

90. Robert, X. & Gouet, P. Deciphering key features in protein structures with the new ENDscript server. Nucleic Acids Res 42, W320–324, doi:10.1093/nar/gku316 (2014).

91. Nicolas, R. H. & Goodwin, G. H. Molecular cloning of polybromo, a nuclear protein containing multiple domains including five bromodomains, a truncated HMG-box, and two repeats of a novel domain. Gene 175, 233–240, doi:10.1016/0378-1119(96)82845-9 (1996).

92. Wang, W. et al. Architectural DNA binding by a high-mobility-group/kinesin-like subunit in mammalian SWI/SNF-related complexes. Proc Natl Acad Sci U S A 95, 492–498, doi:10.1073/pnas.95.2.492 (1998).

93. Satterwhite, E. et al. The BCL11 gene family: involvement of BCL11A in lymphoid malignancies. Blood 98, 3413–3420, doi:10.1182/blood.v98.12.3413 (2001).

